# Glioblastoma Cell Migration is Directed by Electrical Signals

**DOI:** 10.1101/2020.09.09.290254

**Authors:** Hannah Clancy, Michal Pruski, Bing Lang, Jared Ching, Colin D. McCaig

## Abstract

**Background:** Electric field (EF) directed cell migration (electrotaxis) is known to occur in glioblastoma multiforme (GBM) and neural stem cells, with key signaling pathways frequently dysregulated in GBM. One such pathway is EGFR/PI3K/Akt, which is down-regulated by peroxisome proliferator activated receptor gamma (PPARγ) agonists. We investigated the effect of electric fields on GBM differentiated and stem cell migration and whether this was affected by treatment with the PPARγ agonist pioglitazone.

**Methods:** Primary GBM cell lines were cultured as differentiated and glioma stem cells (GSCs) and then exposed to EFs using electrotaxis chambers and imaged with time lapse microscopy. Cells were then treated with varying concentrations of pioglitazone and/or its inhibitor GW9662 and their responses to EFs examined.

**Results:** We demonstrated that GBM differentiated and GSCs have opposing preferences for anodal and cathodal migration, respectively. Pioglitazone treatment resulted in significantly decreased directed cell migration in both cell types. Western blot analysis did not demonstrate any change in PPARγ expression with and without exposure to EF.

**Conclusions:** Opposing EF responses in primary GBM differentiated cells and GSCs can be inhibited chemically by pioglitazone, implicating GBM EF modulation as a potential target in preventing tumour recurrence.

## Introduction

Glioblastoma multiforme (GBM) is the most frequent and aggressive primary brain tumour and is classified by the World Health Organisation as a Grade IV astrocytoma [1,2]. GBM occurs at all ages, however the majority of patients are diagnosed in later life, at a median age of 64 years [2]. Without treatment, the expected survival of those diagnosed with GBM averages 3 months [3], and is extended to 10-15 months with combined surgical resection, chemo- and radiotherapy [4]. Despite these therapeutic advancements and the significant progress in treating systemic cancers, disease recurrence renders GBM incurable. Recurrence is largely attributed to the highly infiltrative nature of GBM, leading to residual cells being inevitably spared by surgical resection [5].

Many of the malignant characteristics of GBM are thought to be attributed to glioma stem cells (GSCs), first isolated in 2002 [6–9]. This subpopulation of GBM cells is considered highly tumourigenic due to the stem-like properties of self-renewal, multilineage differentiation, dysregulated proliferation and increased resistance to apoptosis [10]. The presence of GSCs in GBM has been shown to negatively impact survival [11,12], and is implicated in the invasive behaviour of GBM [5]. Therefore, understanding GSC migration and consequent invasion provides opportunities for targeted therapeutic strategies.

GBM cell migration occurs primarily along white matter tracts and perivascular regions within the brain, through a process whereby the cell becomes morphologically polarised due to underlying cytoskeletal changes [13–15]. Interestingly, neural stem cell (NSC) migration has been shown to be directed by electrical fields (EFs), with cells migrating towards the cathode [16–22]. EFs occur physiologically as a product of ionic and consequently voltage gradients established through spatial variations in ion channels, pumps and leaks [23]. These have been demonstrated in the mammalian brain in vivo, for example along the rostral migratory stream (RMS) [18]. Furthermore, EF guided cell migration; electrotaxis; has been demonstrated in several types of cancer at endogenous voltages, including prostate, breast and more recently brain cancers [24–27]. Limited research has been carried out on the electrotaxis of GBM. Li and colleagues reported cathodal migration in several immortalised glioma cell lines [24], whilst Huang and colleagues demonstrated anodal GSC migration and cathodal migration of differentiated cells [27].

The PI3K/Akt signalling pathway is heavily implicated in cell electrotaxis [17,24,20,26,21,28]. One such mechanism involves a cathodal redistribution of epidermal growth factor receptors (EGFR), resulting in polarised activation of the PI3K/Akt pathway and subsequent polarised actin remodelling [20,29]. This produces asymmetrical membrane protrusions and consequently migration in the direction of the cathode. This pathway is frequently dysregulated in GBM through EGFR overexpression (65% of GBM), and downregulation of the Akt inhibitor PTEN (78% of GBM)[30]. We therefore sought to investigate the effect of an applied EF on the migration of primary GBM cell lines, both differentiated; HROG02-Diff, HROG05-Diff and HROG24-Diff; and de-differentiated, GSC-like; HROG02-GSC, HROG05-GSC with the aim of providing a greater understanding of the infiltrative behaviour of GBM. This is the first time EF migration has been investigated in both differentiated and de-differentiated phenotypes of the same cell lines. Furthermore, we investigated the potential of peroxisome proliferator-activated receptor γ (PPARγ) in modulating the PI3K/Akt pathway.

PPARγ activation has been shown to upregulate PTEN, causing downstream inhibition of Akt [31,32]. PPARγ activation also has been shown to have anti-neoplastic activity on GBM cells [33,34] and anti-proliferative effects on patient derived GSC lines, albeit heterogeneously [35]. The potential role of Akt in electrically guided migration of GBM in conjunction with the high frequency of Akt activating mutations led us to investigate the effect of the PPARγ agonist pioglitazone in both differentiated (HROG02-Diff and HROG05-Diff) and GSC (HROG02-GSC and HROG05-GSC) primary GBM cell lines.

## Materials and Methods

### Reagents

Dulbecco’s Modified Eagle’s Medium, F12 nutrient, penicillin/streptomycin, 0.25% Trypsin-EDTA, Neurobasal-A medium, glutamine, B27, N2 and CO_2_ independent medium were obtained from Invitrogen (Camarillo, CA, USA) and foetal calf serum, EGF, βFGF, heparin, donkey serum, ECM and agar were obtained from Sigma Aldrich (St. Louis, MO, USA). Silicone adhesive (3140 RTV coating) and DC4 high vacuum grease were purchased from Dow Corning (Midland, MI, USA). Steinberg solution was prepared using NaCl and Tris Base sourced from Fisher Bioreagents (USA), KCl, Ca(NO_3_)_2_, and MgSO_4_ obtained from Sigma Aldrich (St. Louis, MO, USA). Primary antibodies for sox-2 (MAB4423) and nestin (ABD69) were purchased from Millipore (USA), whilst primary antibodies for GFAP (G3893) were obtained from Sigma Aldrich (St. Louis, MO, USA), CD133 from Miltenyi Biotec (Germany). Alexa Fluor 594 donkey anti-mouse (R37115), Alexa Fluor 594 donkey anti-rabbit (R37119), Alexa Fluor 488 donkey anti-goat (A-11055) and Alexa Fluor 488 donkey antimouse (A-21202) were purchased from Molecular Probes (USA). Pioglitazone hydrochloride (E6910), GW9662 (M6493) and phenozine methosulfate was purchased from Sigma Aldrich (St. Louis, MO, USA) and XTT cell viability assay from Invitrogen (USA). Stock solutions of 10000μM pioglitazone:DMSO and 10000μM GW9662:DMSO were prepared. Lysis buffer solution was prepared using cell lytic lysis buffer obtained from Sigma Aldrich (St. Louis, MO, USA) and complete EDTA free and phosphostop protease inhibitors from Roche, (Switzerland). Western blotting was completed using 4-12% Bis-Tris pre-cast gels and MOPS running buffer purchased from Invitrogen (USA), Nitrocellulose membrane obtained from GE Healthcare (Amersham, United Kingdom), Anti-PPARγ (E-8) (sc-7273) purchased from Santa Cruz, (USA), Anti-β-Actin from Sigma Aldrich (A2228) (St. Louis, MO, USA) and 680 Alexa Fluor donkey anti-mouse IgG (A-10038) secondary antibodies purchased from Molecular Probes (USA).

### Cell Culture

Primary GBM Cell lines, HROG02, HROG05 and HROG24, were kindly provided by Professor Linnebacher of the University of Rostock. To produce the differentiated phenotype, cells were maintained in DMEM:F12 supplemented with 5% foetal calf serum and 1% penicillin/streptomycin at 37°C, 5% carbon dioxide. This produced three differentiated cell lines: HROG02-Diff, HROG05-Diff and HROG24-Diff. To achieve the de-differentiated, GSC phenotype, cells were cultured in Neurobasal-A medium supplemented with 2mM glutamine, 1% penicillin/streptomycin, epidermal growth factor (20ng/mL), basic fibroblast growth factor (20ng/mL), heparin (2μg/mL), 2% B27 and 1% N2 at 37°C, 5% carbon dioxide [36–38]. De-differentiated cells formed neurospheres when seeded in ultra-low adhesion 6 well plates and were termed HROG02-GSC and HROG05-GSC.

### Electrotaxis Assay

Assays to assess cell electrotaxis were adapted from a previously described protocol [39]. Electrotactic chambers were constructed by gluing No. 1 thickness cover glass slides to the base of a 10cm petri-dish using silicone adhesive: the resulting electrotactic region measured 10 x 40mm. Cells were seeded on top of a layer of 1%ECM within the electrotactic region and incubated at 37°C, 5% carbon dioxide overnight. High vacuum grease was used to apply a No. 1 thickness glass slide roof over the electrotactic region shortly before experimentation, and 5ml CO2 independent medium added (CO_2_ independent media replaced DMEM:F12 and Neurobasal-A medium in differentiated and GSC cultures, respectively). EFs were delivered via silver/silver chloride electrodes in reservoirs of Steinberg’s solution, passing current through 2% Agar bridges into the insulated pools of CO_2_ independent medium on the left (cathode) and the right (anode) side of the electrotactic chamber. Cells were exposed to either 0 (control), 50, 100 or 200mV/mm for 3 hours, with images taken every 10 minutes using a Leica DM IRB inverted microscope, digital camera and Velocity software (Improvision, UK).

### Quantification of Electrotaxis

Image J software (National Institution of Health, USA), was used to track cells manually, and Microscoft Excel used to calculate velocity (total distance/time) and directedness. Directedness was calculated by Cosine(θ) where θ is the angle formed between the cell trajectory and EF vector. This gives a value between +1 (perfect anodal migration) and −1 (perfect cathodal migration), with migration perpendicular to the EF vector producing a value of 0. Experiments were repeated three times, with 100 cells tracked per experiment, n=300 for all velocity and directedness data. The Chemotaxis plugin (Ibidi GmbH, Germany) for Image J was used to produce migration diagrams, animations and data for xFMI (proportion of cell migration in the x-axis) change over time. The number of cells migrating to either the anode or cathode was obtained from migration diagrams, resulting in three counts of migration for both directions (n=3 for cell count data).

### Immunocytochemistry

Immunocytochemistry was completed concurrently on differentiated and corresponding dedifferentiated primary GBM cell lines. Cells were seeded onto 1% ECM coated circular borosilicate cover slips and incubated at 37°C, 5% carbon dioxide overnight. Slides were fixed with 4% paraformaldehyde and permeabilised with 0.3% Triton-X-100. Cells were stained with primary antibodies for sox-2 (1:200), CD133 (1:50), nestin (1:500) and GFAP (1:400) overnight at 4°C. Slides were probed with Alexa Fluor fluorescent secondary antibodies and counterstained with Hoechst (1:2000). For fluorescent microscopy, a Zeiss imager M2 upright microscope with DAPI (Zeis set 49), DsRed (Zeis set 43) and FITC (Zeiss set 10) filters was used, with images captured using a high resolution microscope camera and Axiovision software (for all images except for HROG05-Diff and HROG05-GSC stained for Hoechst, GFAP and nestin). These later images were captured using a high resolution camera linked to a Zeiss Axio Observer Z1 inverted microscope using DAPI (Zeiss set 49), Texas Red (Semrock) and FITC (Zeiss set 10) filters, with axiovision software (Germany).

CD133, nestin and sox-2 positive cells are associated with the GSC phenotype, whilst GFAP staining is astrocyte specific and therefore indicates a differentiated phenotype [37,40].

### Drug Treatments

XTT viability assays were performed in triplicate, testing different concentrations of the PPARγ agonist pioglitazone and PPARγ antagonist GW9662. Cells were exposed to treatments for 24, 48 or 72 hours at 37°C, 5% carbon dioxide, after which XTT/PMS solution was added and the absorbance of the individual wells read at 450nm. As the XTT viability assay omitted the application of an EF, results were used to guide trial electrotaxis assays. This lead to the experimental condition of 15μM pioglitazone + 5μM DMSO (pioglitazone treatment) and control conditions of 15μM pioglitazone + 5μM GW9662 (piogliazone/GW9662 treatment) and 20μM DMSO (DMSO treatment), exposed for between 12-24 hours. Electrotaxis assays were completed as previously described. The cells were seeded into the electrotaxis chambers in media containing drug treatments, with the time of application noted. Upon experimentation, CO2 independent media containing drug treatments was added and an EF of 200mV/mm was applied for 3 hours. Total time of treatment exposure was recorded; there were no significant differences in total exposure times.

### Western Blotting

Collection of protein samples was achieved using the method as described for electotaxis assay, however using a larger 40 x 60mm electrotactic region. This larger region resulted in increased voltage instability and temperature gain over a 3 hours period. Preliminary data concerning xFMI change with time showed that xFMI generally peaked after one hour of exposure to 200mV/mm EF, therefore this time period and field strength was applied to samples. This was repeated three times for EF completed simultaneously with a paired control condition, following an identical procedure but with no EF applied. Adherent cells were lysed using lysis buffer solution and separated by electrophoresis on 4-12% Bis-Tris pre-cast gels using MOPS running buffer and transferred onto nitrocellulose membrane. Membranes were cut and incubated with appropriate primary antibody; anti-PPARγ (1:100) or β-actin (1:50000); and subsequently 680 donkey anti-mouse secondary antibody. Membranes were visualised by infrared scanner (Li-Cor Odyssey, USA) and quantified using Odyssey 2.1 software (USA) to produce an Integrated Intensity value (II). PPARγ II was normalised to β-actin control.

#### Statistics

All graphs and statistical analysis was performed using Graphpad Prism 5.04, (USA). All migration data was analysed by One-Way ANOVA and Tukey’s multiple comparison post-test, with *p*<0.05 considered significant. Western blot data was tested using Unpaired T-test, deemed significant at *p*<0.5. All data presented as mean±SEM. Results from Tukey’s post-test were given as follows: Ns = non-significant,* = significant (0.01<*p*<0.05), ** = very significant (0.001<*p*<0.01), *** = extremely significant (*p*<0.001).

### Results

#### HROG02-GSC and HROG05-GSC express glioma stem cell markers

HROG02-Diff stained negatively for sox-2, CD133, weakly for nestin and positively for GFAP (Fig. 1a). In contrast, HROG02-GSC stained strongly for the GSC markers sox-2, CD133 and nestin, whilst demonstrating variable GFAP expression (Fig. 1b). This staining pattern is in keeping with the distinct differentiated and GSC phenotypes.

**Figure 1.**
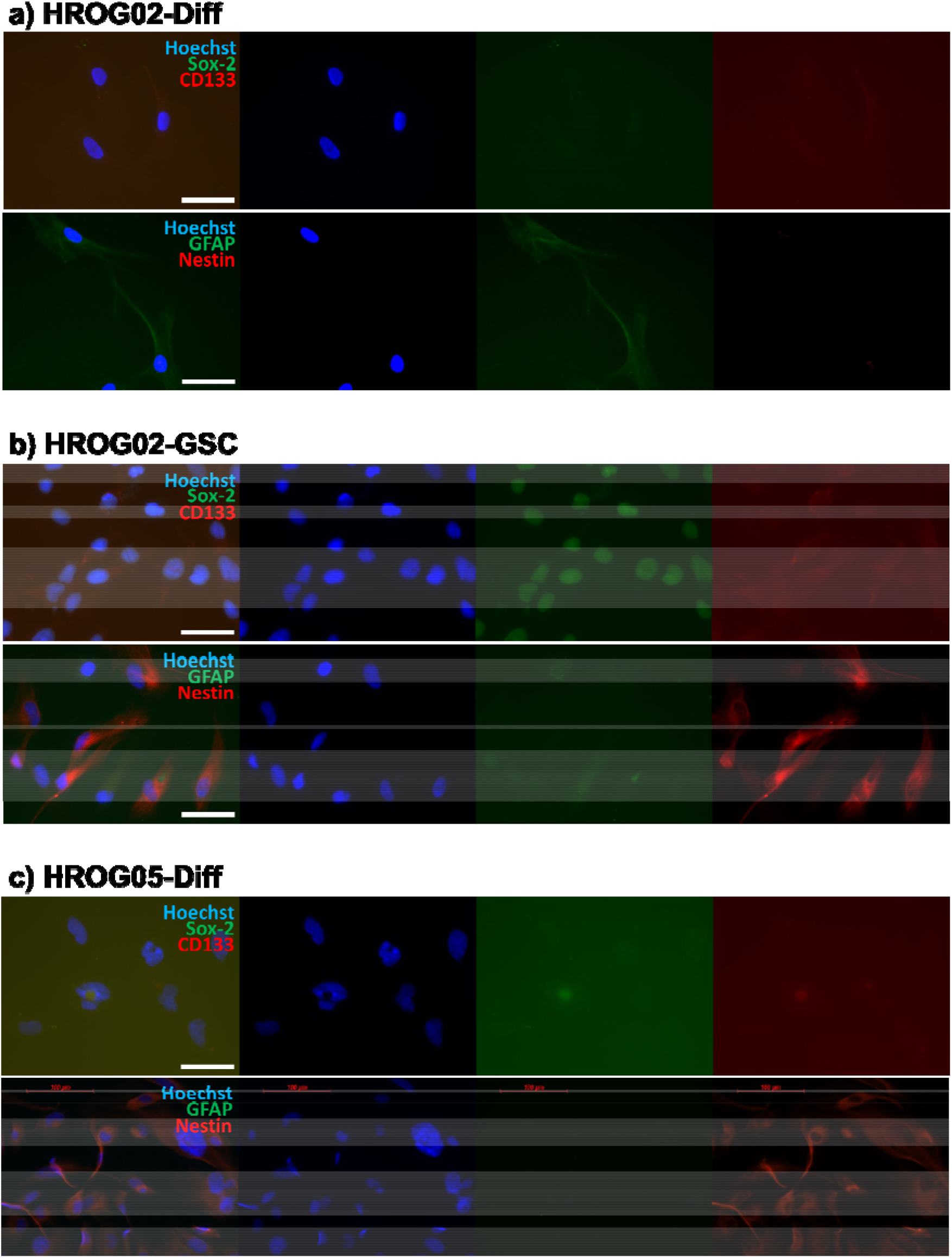
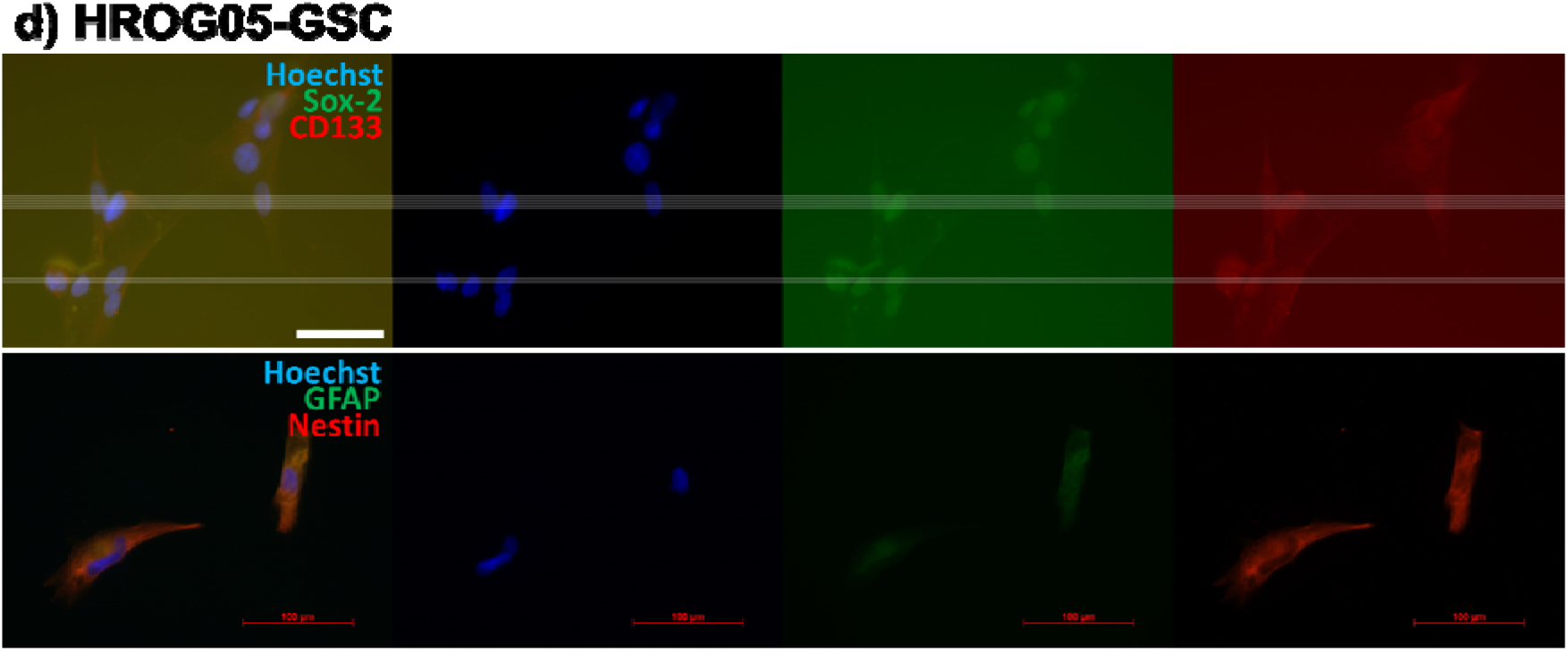
Immunocytochemistry staining of HROG02-Diff (a), HROG02-GSC (b), HROG05-Diff (c), and HROG05-GSC (d). Nuclei are visualised in blue (Hoechst staining), green visualised either ant-Sox-2 or anti-GFAP, red visualises either anti-CD133 or anti-Nestin. Scale bar represents 100 μm. HROG02-Diff and HROG05-Diff showed negative staining for sox-2 and CD133. HROG02-GSC and HROG05-GSC stained positively for sox-2, CD133 and nestin. GFAP was the marker that showed the most variability in staining.

HROG05-Diff stained negatively for sox-2, CD133 and GFAP, but positively for nestin (Fig. 1c). HROG05-GSC in comparison stained positively for sox-2, CD133, nestin and GFAP (Fig. 1d), however again GFAP showed variable expression across repeated imaging.

#### Primary differentiated cell lines migrate anodally in a voltage dependent manner

Figures 2a, 3a and 4a demonstrate a higher proportion of HROG02-Diff, HROG05-Diff and HROG24-Diff cells migrated towards the anode when exposed to a greater EF. Figures 2d, 3d and 4d show that with no EF cells did not migrate preferentially to either pole. However, application of EF induced predominantly anodal migration of cells in a voltage dependent manner, with 84.3±4.1% (HROG02-Diff), 71.0±7.0% (HROG05-Diff) and 89.7±0.9% (HROG24-Diff) of cells migrating towards the anode at 200mV/mm. Figures 2b, 3b and 4b demonstrate a stepwise increase in directedness of cell migration towards the anode in all three differentiated cell lines, with statistically significant changes in the presence of an EF as low as 50mV/mm in HROG02-Diff and HROG24-Diff, and 100mV/mm in HROG05-Diff. Increasing EF was also associated with increasing velocity in cell migration, as shown in Figures 2c, 3c and 4c. EFs as low as 50mV/mm produced statistically significant increases in velocity compared to control; 6.6±0.5μm/hour (no EF) to 11.7±0.5μm/hour at 50mV/mm (HROG02-Diff), 6.5±0.4 to 10.7±0.6μm/hour (HROG05-Diff), and 5.0±0.4 to 9.3±0.4μm/hour (HROG24-Diff). Figure 2e, 3e and 4e show the change in xFMI over the duration of the experiment, with all three differentiated cell lines showing a plateau in the proportion of migration towards the anode by approximately 60 minutes.

**Figure 2.**
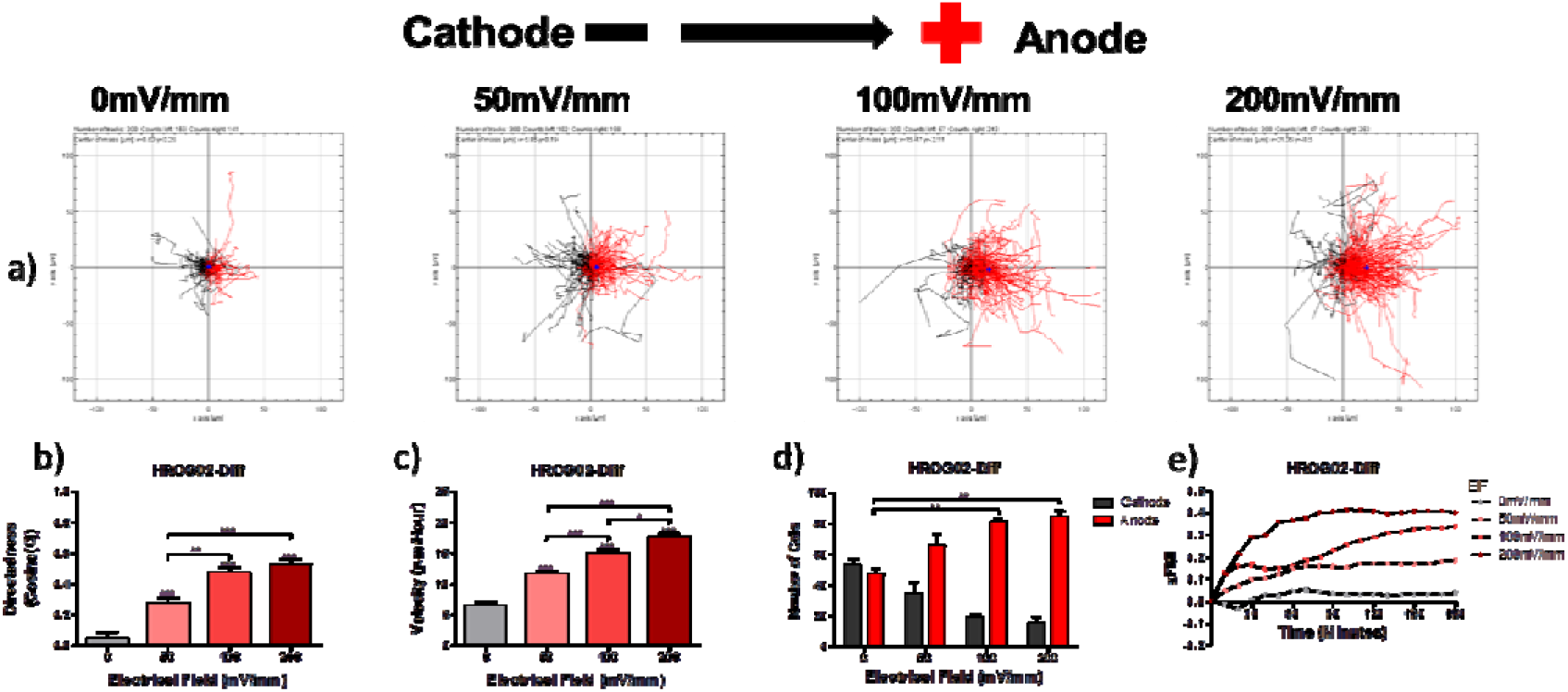
HROG02-Diff cell migration in electric fields. Panel (a) visualises the cellular movement that occurred under different field strengths. (b-d) show, respectively, the effects of field strength on the directness of migration, cell velocity, and the number of cells that migrated towards each electrode. (e) illustrates the effect of field strength over time on the x-Forward Migration Index. Cells show an anodal migration pattern (a; p=0.0022 in d), with a statistically significant effect of the electric field on directness (b, p<0.0001) and velocity (c, p<0.0001); one-way ANOVA. *, ** and *** above bars represent Tukey’s post-test comparisons with control (0 mV/mm), while horizontal bars represent Tukey’s results of comparisons between groups that have shown statistical significance.

**Figure 3.**
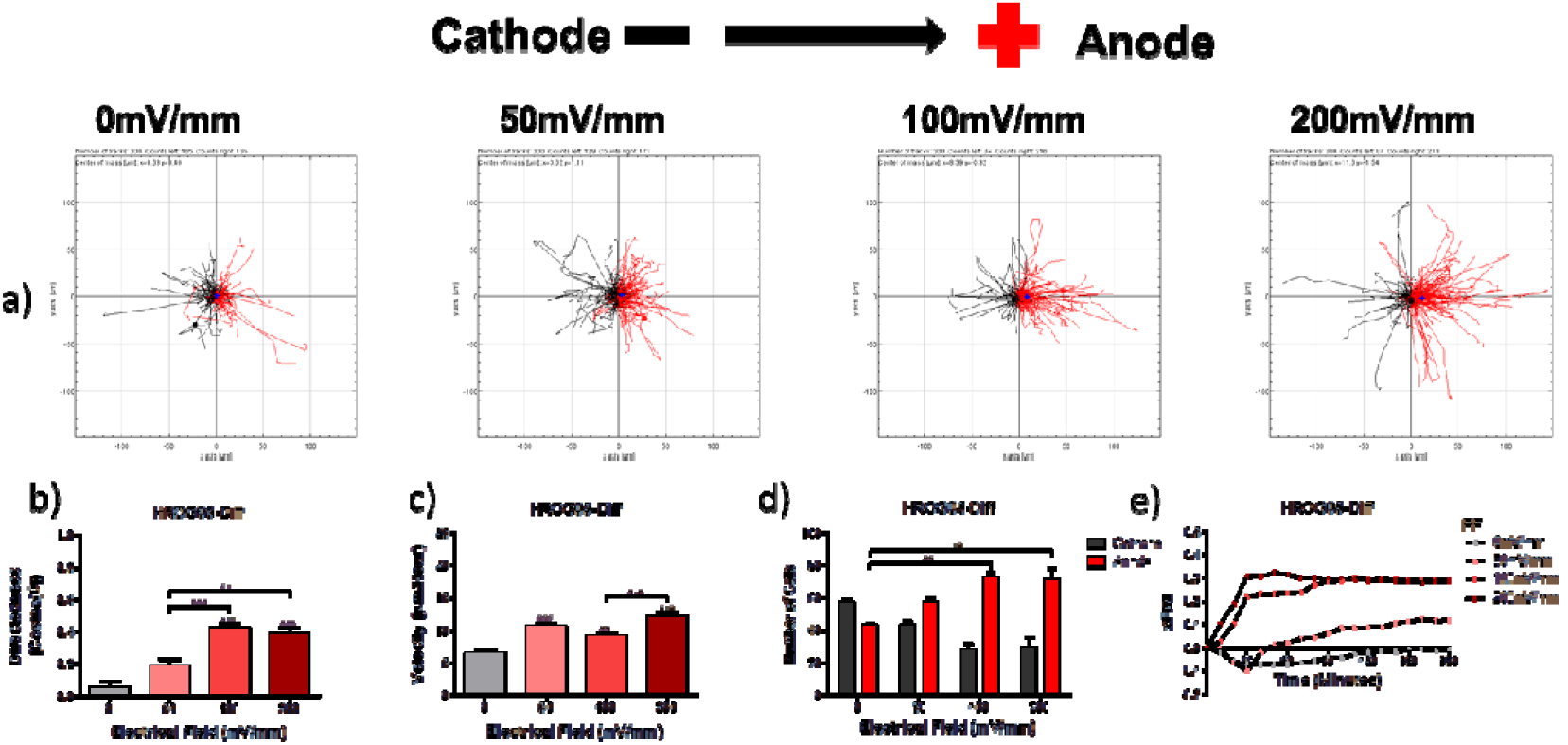
HROG05-Diff cell migration in the electric fields. Panel (a) visualises the cellular movement that occurred under different field strengths. (b-d) show, respectively, the effects of field strength on the directness of migration, cell velocity, and the number of cells that migrated towards each electrode. (e) illustrates the effect of field strength over time on the x-Forward Migration Index. Cells show an anodal migration pattern (a; p=0.0035 in d) with a statistically significant effect of the electric field on directness (b, p<0.0001) and velocity (c, p<0.0001); one-way ANOVA. *, ** and *** above bars represent Tukey’s post-test comparisons with control (0 mV/mm), while horizontal bars represent Tukey’s results of comparisons between groups that have shown statistical significance.

**Figure 4.**
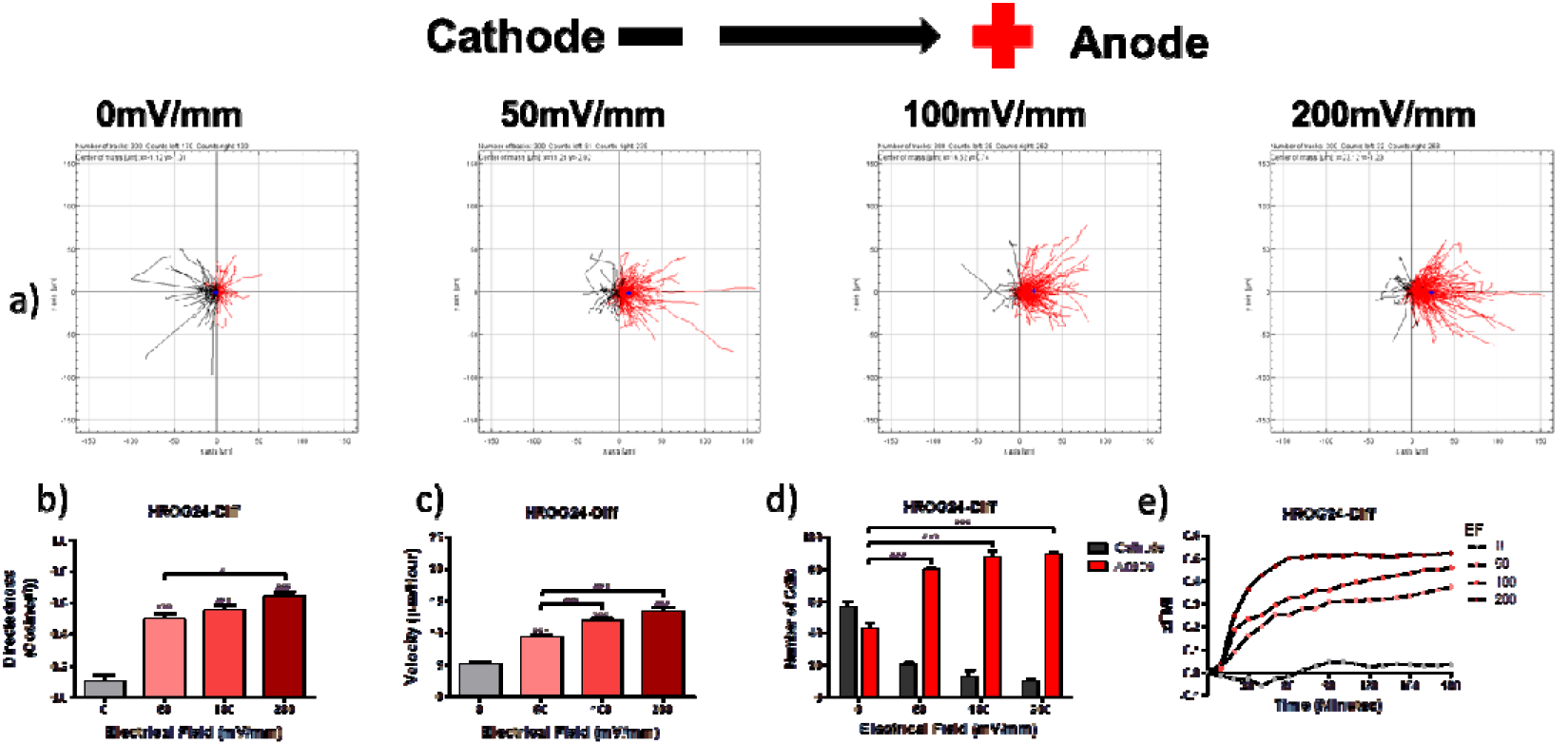
HROG24-Diff cell migration in electric fields. Panel (a) visualises the cellular movement that occurred under different field strengths. (b-d) show, respectively, the effects of field strength on the directness of migration, cell velocity, and the number of cells that migrated towards each electrode. (e) illustrates the effect of field strength over time on the x-Forward Migration Index. Cells show an anodal migration pattern (a; p<0.0001 in d) with a statistically significant effect of the electric field on directness (b, p<0.0001) and velocity (c, p<0.0001); one-way ANOVA. *, ** and *** above bars represent Tukey’s post-test comparisons with control (0 mV/mm), while horizontal bars represent Tukey’s results of comparisons between groups that have shown statistical significance.

#### EF induces cathodal migration in GSC cell lines

In contrast to primary differentiated cell lines, application of increasing EF resulted in a greater proportion of GSC line cells migrating towards the cathode (Fig. 5a and 6a). Increasing the EF applied caused a corresponding increase in the percentage of cells migrating towards the cathode, increasing from 49.3±2.9% of cells with no EF, to 70.3±2.7% cells with an EF of 200mV/mm in HROG02-GSC, and from 44.7±5.2% cells with no EF present, to 68.3±3.7% cells at 200mV/mm in HROG05-GSC (Fig. 5d and 6d). This was further reflected in directedness, which increased towards the cathode in the presence of an EF (Fig. 5b and 6b). Application of 50mV/mm to HROG02-GSC produced a statistically significant increase in directedness towards the cathode compared to control from 0.15±0.04 to −0.12±0.04, remaining at similar directedness when exposed to 100mV/mm (−0.11±0.04), before significantly increasing to −0.28±0.04 at 200mV/mm (Fig. 5b). Application of an EF as low as 100mV/mm (−0.25±0.03) induced a significant increase in directedness of HROG05-GSC cell migration towards the cathode when compared to no EF applied (0.17±0.04) (Fig. 6b). Increasing the EF applied to 200mV/mm did not significantly increase directedness compared to 100mV/mm. HROG02-GSC showed increased velocity of cell migration with increased field strength (Fig. 5c); mean cell velocity when exposed to 200mV/mm was approximately 60% higher than when no EF was applied. With increasing EF applied to HROG05-GSC cells, velocity initially decreased compared to control in an EF of 50mV/mm (Fig. 6c). However, subsequent increases in EF strength was correlated with greater velocity of cell migration. Figures 5e and 6e show the change in xFMI over time in HROG02-GSC and HROG05-GSC respectively, demonstrating a plateau in xFMI reached by 60 minutes. Upon exposure to an EF of 100 and 200mV/mm, HROG05-GSCs initially appear to migrate towards the anode, a positive xFMI, in a similar manner to the control group, however by 30 minutes began to steadily migrate more cathodally (Fig. 6e).

**Figure 5.**
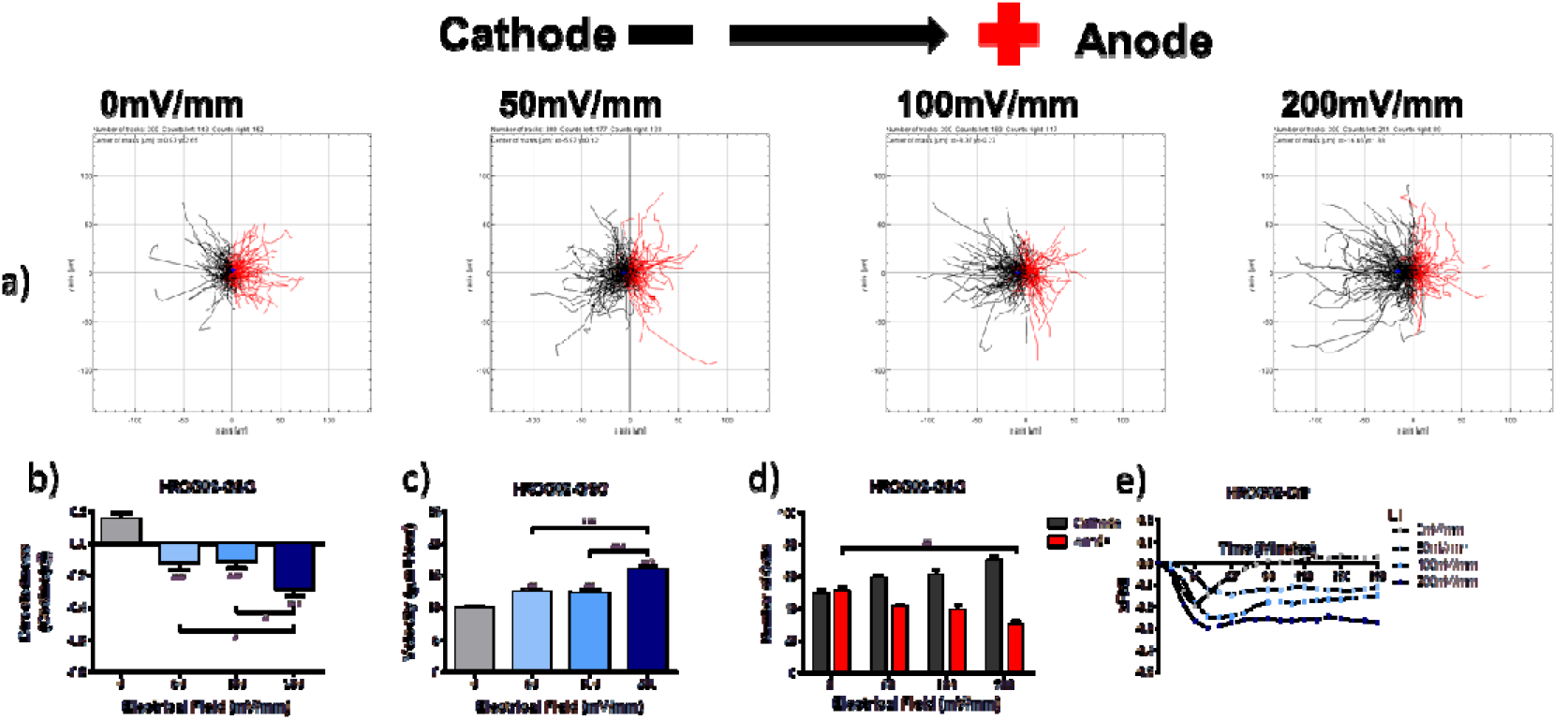
HROG02-GSC cell migration in electric fields. Panel (a) visualises the cellular movement that occurred under different field strengths. (b-d) show, respectively, the effects of field strength on the directness of migration, cell velocity, and the number of cells that migrated towards each electrode. (e) illustrates the effect of field strength over time on the x-Forward Migration Index. Cells show a cathodal migration pattern (a; p=0.0034 in d) with a statistically significant effect of the electric field on directness (b, p<0.0001) and velocity (c, p<0.0001); one-way ANOVA. *, ** and *** above bars represent Tukey’s post-test comparisons with control (0 mV/mm), while horizontal bars represent Tukey’s results of comparisons between groups that have shown statistical significance.

**Figure 6.**
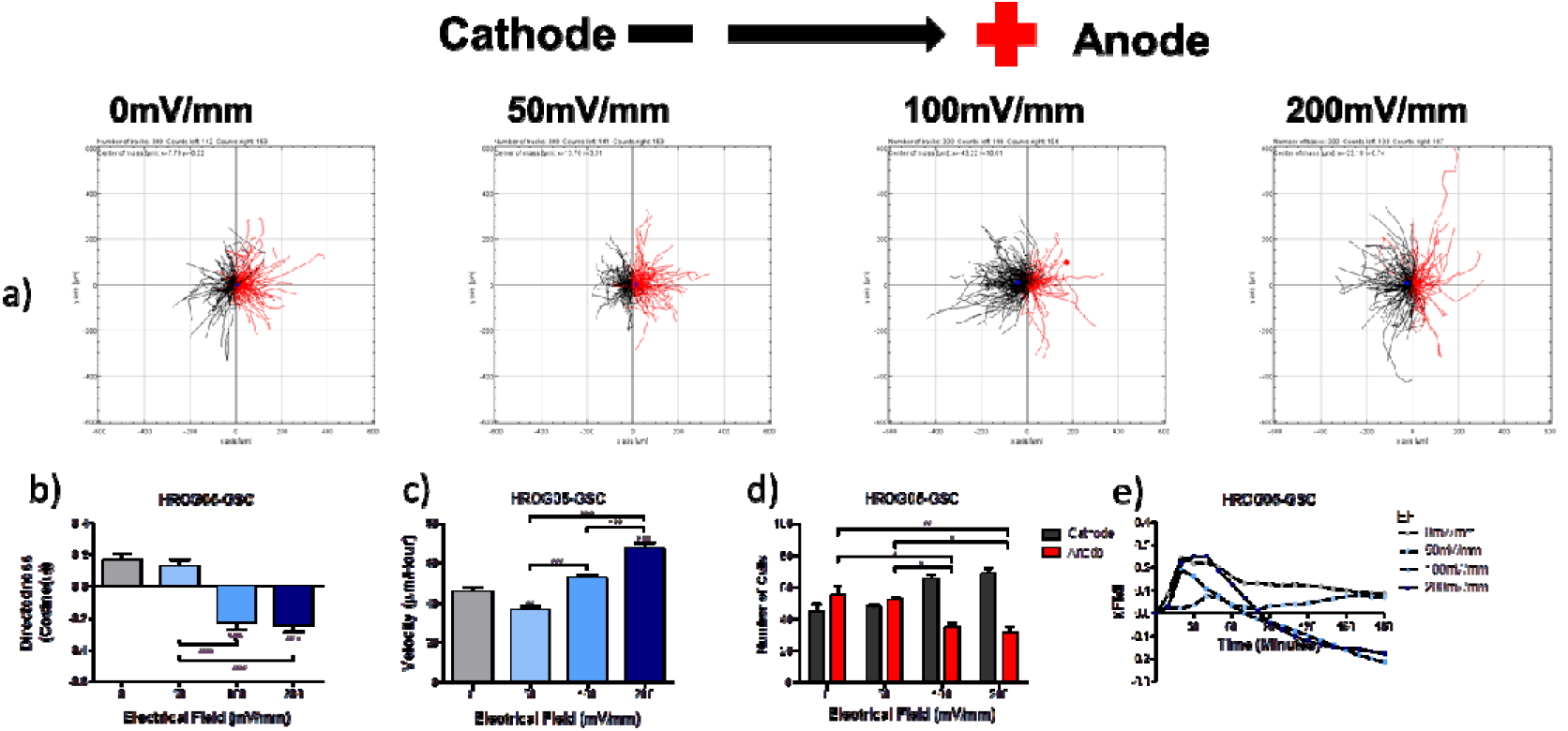
HROG05-GSC cell migration in electric fields. Panel (a) visualises the cellular movement that occurred under different field strengths. (b-d) show, respectively, the effects of field strength on the directness of migration, cell velocity, and the number of cells that migrated towards each electrode. (e) illustrates the effect of field strength over time on the x-Forward Migration Index. Cells show a cathodal migration pattern (a; p=0.0030 in d) with a statistically significant effect of the electric field on directness (b, p<0.0001) and velocity (c, p<0.0001); one-way ANOVA. *, ** and *** above bars represent Tukey’s post-test comparisons with control (0 mV/mm), while horizontal bars represent Tukey’s results of comparisons between groups that have shown statistical significance.

#### Pioglitazone significantly alters EF guided migration of HROG05-Diff and HROG02-Diff primary GBM cell lines

Application of differing pharmacological treatments induced a statistically significant change in the directedness of cell migration of HROG02-Diff and HROG05-Diff, *p*=0.0280 and *p*<0.0001 calculated respectively by One-Way ANOVA (Fig. 7b, 8b). Directedness of HROG02-Diff cell migration towards the anode decreased to 0.28±0.04 upon exposure to pioglitazone treatment; a decrease of approximately 30% compared to both DMSO and pioglitazone/GW9662 treatments. The percentage of HROG02-Diff cells migrating towards the anode decreased with pioglitazone treatment (67.0±1.5%) compared to DMSO (76.7±4.7%) and pioglitazone/GW9662 (78.7±4.2%) (Fig. 7d). There was no significant change in HROG02-Diff cell migration velocity between treatment conditions (*p*=0.3592) (Fig. 7c). HROG05-Diff cells also showed a statistically significant decrease in anodal migration when treated with pioglitazone compared to both DMSO and pioglitazone/GW9662 treatments. This was represented as both a significant decrease in the directedness of anodal migration (Fig. 8b) and in the number of cells migrating towards the anode (Fig. 8d). Directedness with pioglitazone treatment was 0.10±0.04, whilst 0.36±0.04 with DMSO and 0.39±0.04 with pioglitazone/GW9662; a statistically significant decrease (Fig. 8b). Pioglitazone treatment also induced a significant decrease in HROG05-Diff cell velocity (Fig. 8c). Figure 7e shows a similar pattern of xFMI change over time of HROG02-Diff between the three treatment conditions, however, after approximately 60 minutes, the xFMI of cells treated with pioglitazone slowly decreased. HROG05-Diff cells treated with pioglitazone showed minimal change in xFMI, reaching a peak of 0.05, compared to peaks of 0.34 and 0.35 with DMSO and pioglitazone/GW9662 treatments respectively (Fig. 8e).

**Figure 7.**
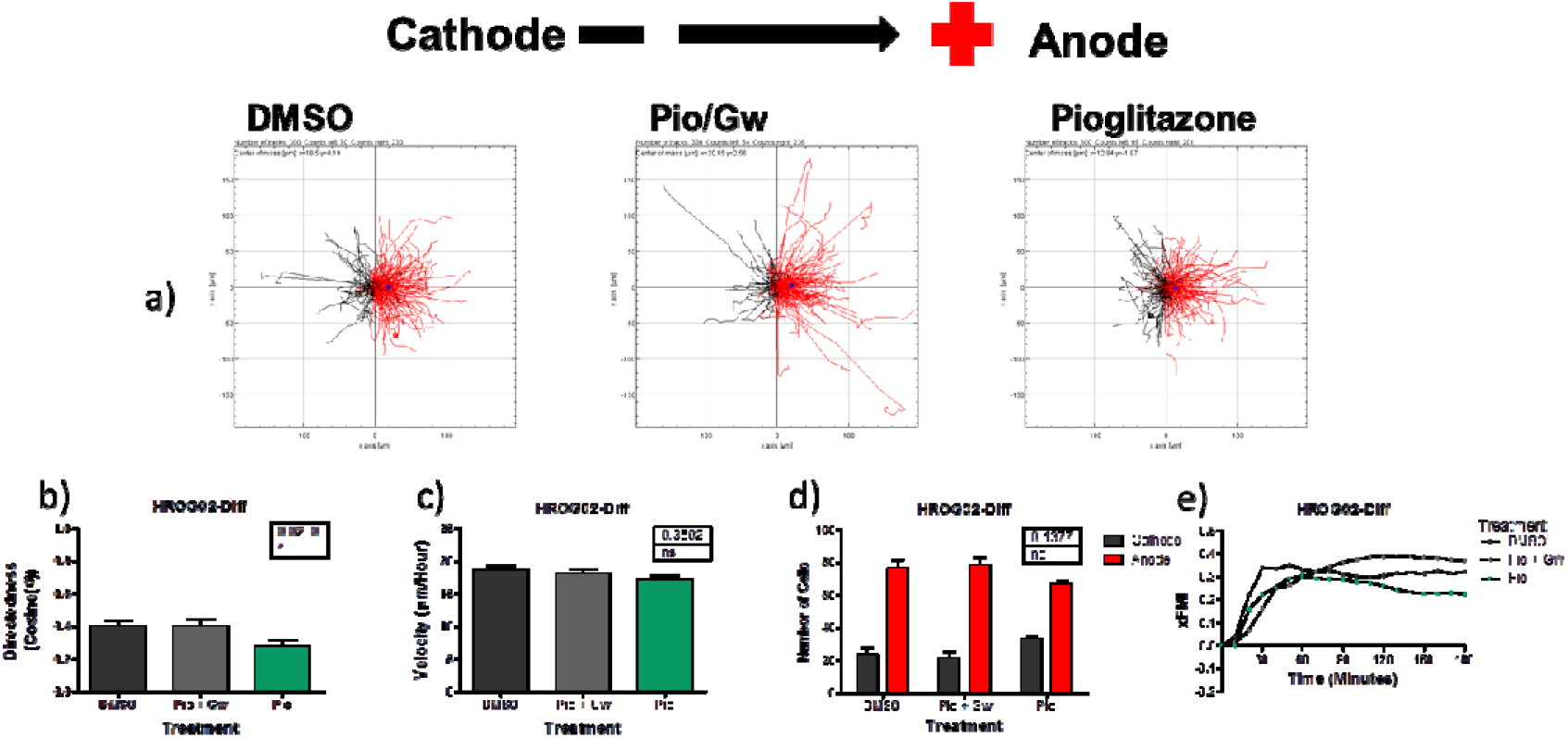
Effect of PPARγ stimulation on HROG02-Diff cell migration in a 200mV/mm electric field. (a) illustrates changes caused by the addition of DMSO (drug vehicle) and PPARγ agonists (pioglitazone) and antagonist (GW9662). (b-d) show, respectively, the effects of field strength on the directness of migration, cell velocity, and the number of cells that migrated towards each electrode. (e) illustrates the effect of treatment over time on the x-Forward Migration Index. Treatment had a significant effect on the cells’ directedness (b, p=0.0280; one-way ANOVA), but Tukey’s post-hoc analysis revealed no significant difference between groups.

**Figure 8.**
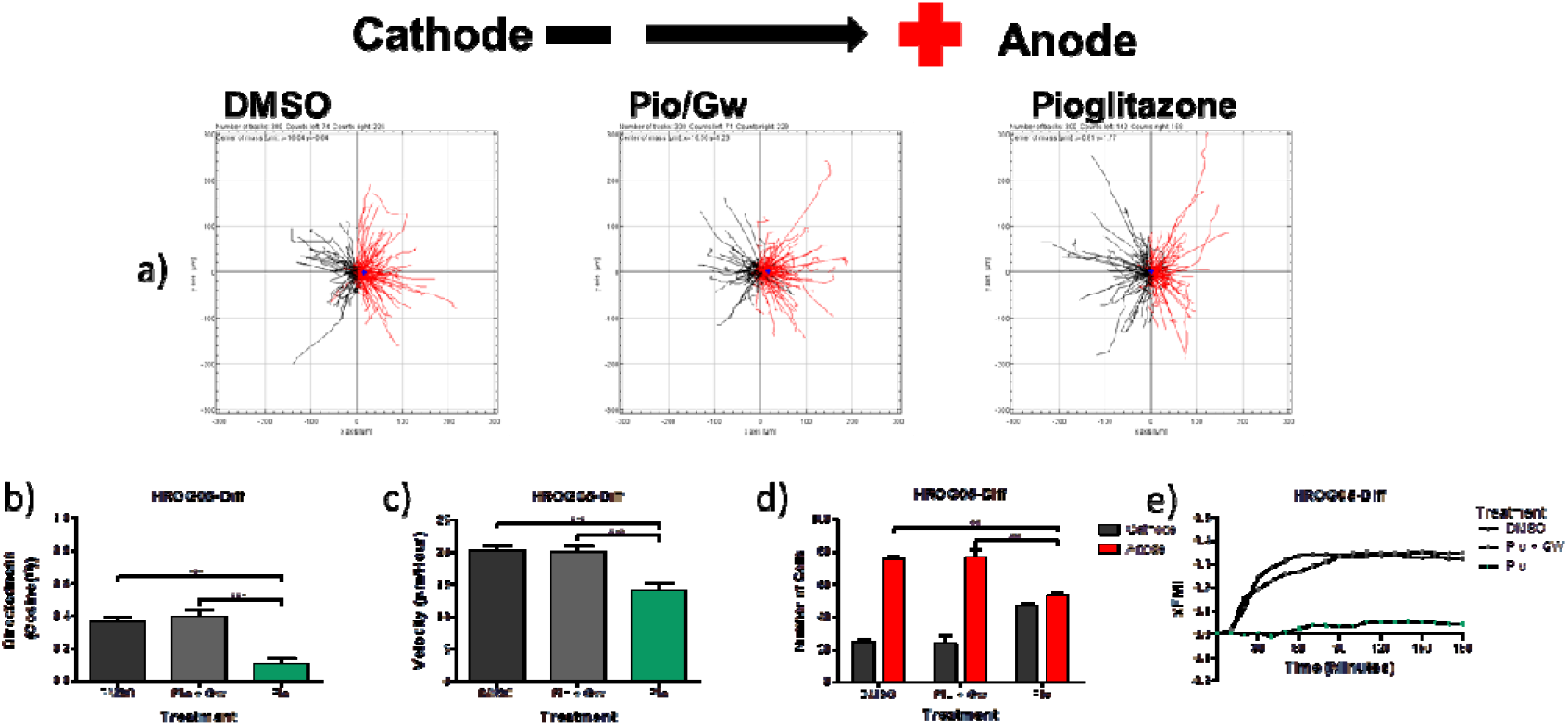
Effect of PPARγ stimulation on HROG05-Diff cell cell migration in a 200mV/mm electric field. (a) illustrates changes caused by the addition of DMSO (drug vehicle) and PPARγ agonists (pioglitazone) and antagonist (GW9662). (b-d) show, respectively, the effects of field strength on the directness of migration, cell velocity, and the number of cells that migrated towards each electrode. (e) illustrates the effect of treatment over time on the x-Forward Migration Index. Treatment had a significant effect on the direction of migration (a; P=0.0034 in d), as well as on directedness (b, p<0.0001) and velocity (c, p<0.0001); one-way ANOVA. *, ** and *** above horizontal bars represent Tukey’s results of comparisons between groups that have shown statistical significance.

#### Pioglitazone significantly decreases directedness of cell migration in GSC cell lines

Figure 9b shows treatment of HROG02-GSC cells with pioglitazone significantly decreased directedness of cell migration towards the cathode compared to both DMSO and pioglitazone/GW9662 treatments. Figure 9a and 9d show that fewer cells migrated towards the cathode when treated with pioglitazone, however this did not represent a statistically significant change. Velocity of cell migration was unaffected by different drug conditions (*p*=0.4484; Fig. 9c). Pioglitazone treatment alone diminished cathodal migration of HROG05-GSC cells demonstrated by a reversal in directional preference (Fig. 10a and 10d), and in directedness of cell migration (Fig. 10b). Figure 10b shows both DMSO (−0.23±0.04) and pioglitazone/GW9662 (−0.38±0.04) treatments produced cathodal migration, however pioglitazone was associated with minimal anodal migration (0.03±0.04); a statistically significant correlation. Velocity of HROG05-GSC cell migration was significantly reduced when treated with pioglitazone compared to pioglitazone/GW9662 however there was no significant decrease in comparison to DMSO treatment (Fig. 10c).

**Figure 9.**
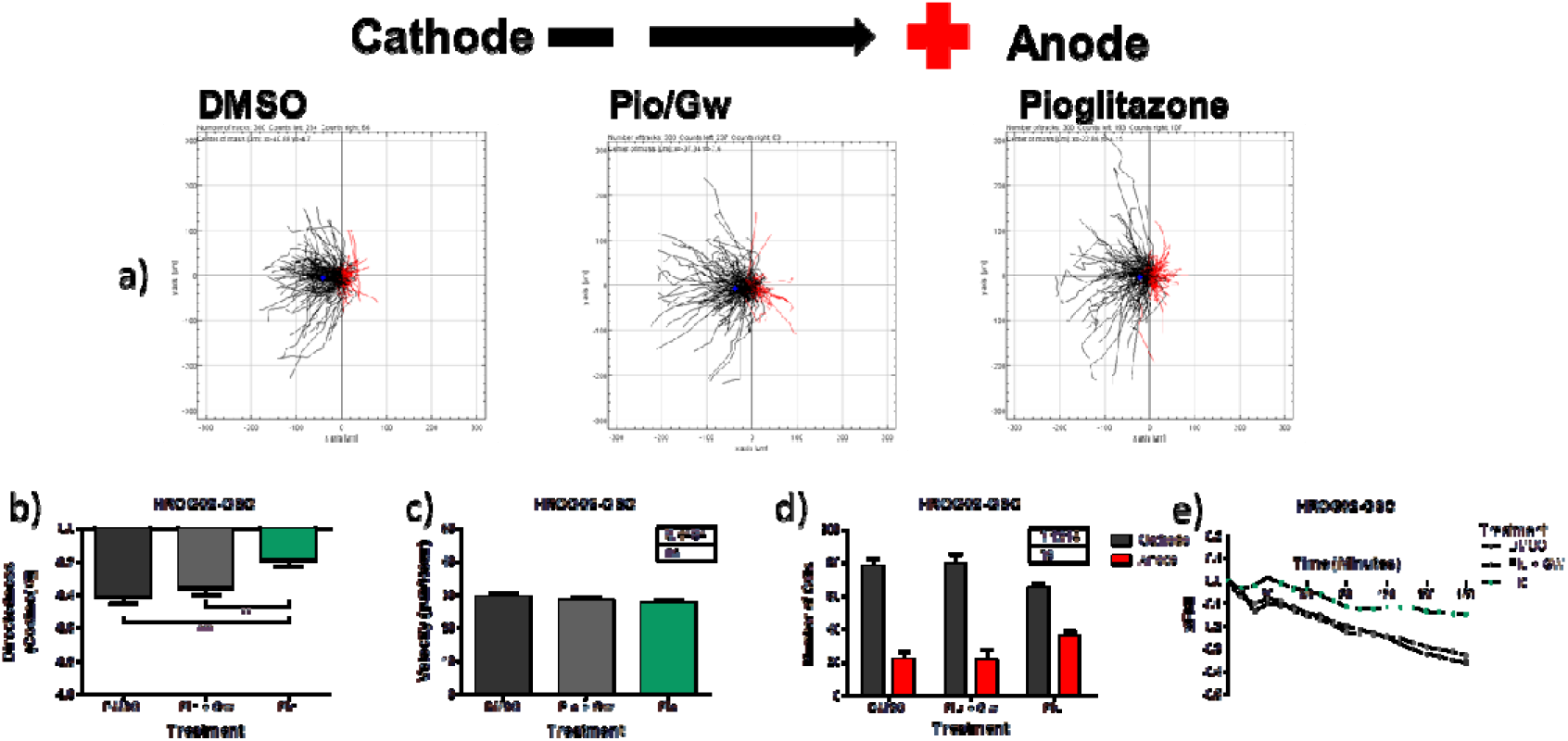
Effect of PPARγ stimulation on HROG02-GSC cell migration in a 200mV/mm electric field. (a) illustrates changes caused by the addition of DMSO (drug vehicle) and PPARγ agonists (pioglitazone) and antagonist (GW9662). (b-d) show, respectively, the effects of field strength on the directness of migration, cell velocity, and the number of cells that migrated towards each electrode. (e) illustrates the effect of treatment over time on the x-Forward Migration Index. Treatment had a significant effect on directedness (b, p=0.0002; one-way ANOVA). *, ** and *** above horizontal bars represent Tukey’s results of comparisons between groups that have shown statistical significance.

**Figure 10.**
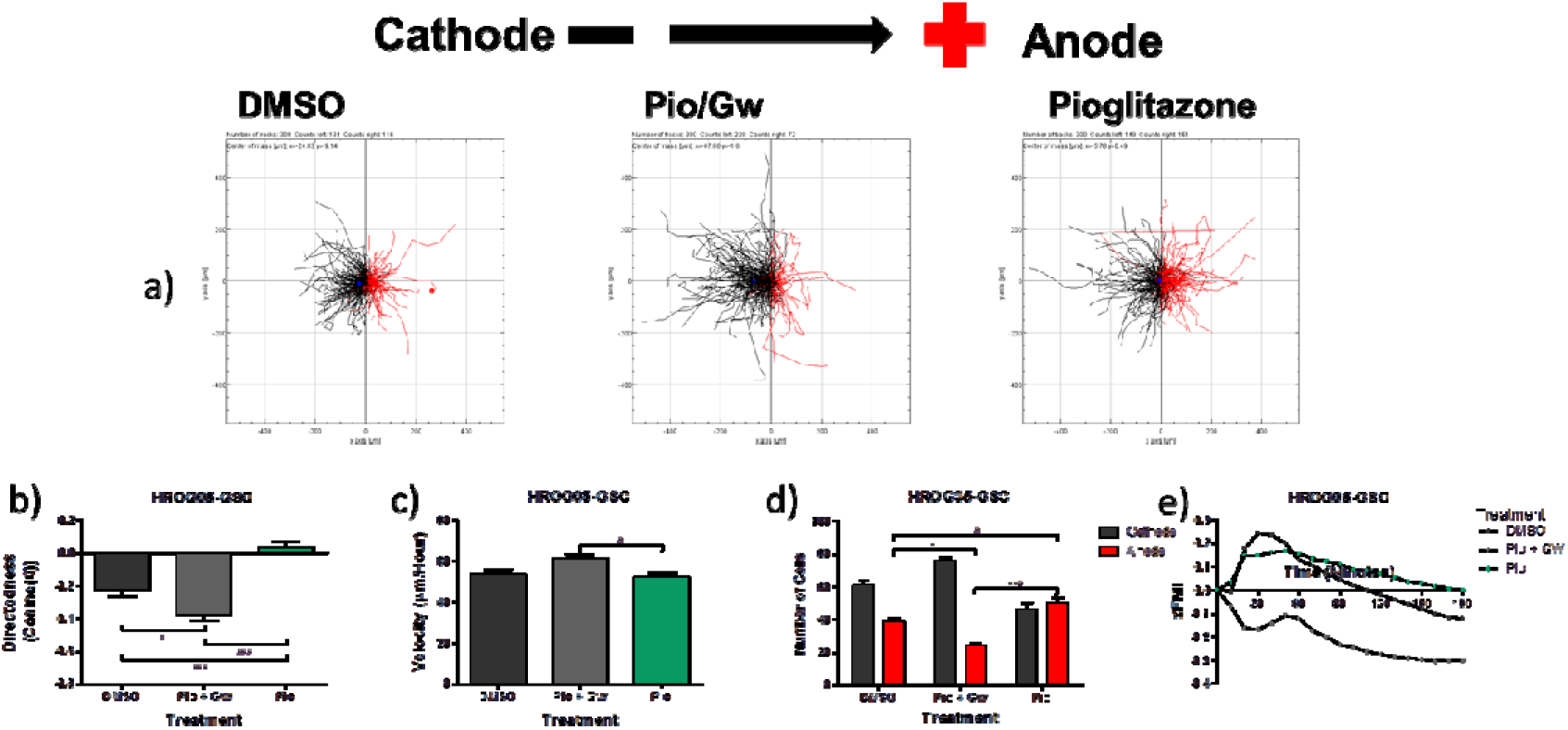
Effect of PPARγ stimulation on HROG05-GSC cell migration in a 200mV/mm electric field. (a) illustrates changes caused by the addition of DMSO (drug vehicle) and PPARγ agonists (pioglitazone) and antagonist (GW9662). (b-d) show, respectively, the effects of field strength on the directness of migration, cell velocity, and the number of cells that migrated towards each electrode. (e) illustrates the effect of treatment over time on the x-Forward Migration Index. Treatment had a significant effect on the direction of migration (a; P=0.0008 in d; one-way ANOVA), though differences between the DMSO, and pioglitazone with GW9662 should be noted. Treatment had also effects on directedness (b, p<0.0001) and velocity (c, p<0.0107); one-way ANOVA. *, ** and *** above horizontal bars represent Tukey’s results of comparisons between groups that have shown statistical significance.

#### EF does not alter PPARγ expression

Figure 11b, 12b, 13b and 14b show infrared scans of western blots stained for PPARγ and β-actin expression in HROG02-Diff, HROG05-Diff, HROG02-GSC and HROG05-GSC cells respectively with and without EF application. Quantification of these reveal no statistically significant change in the normalised Integrated Intensity of PPARγ in any cell lines tested upon application of an EF (*p*=>0.5) (Fig. 11a, 12a, 13a and 14a).

**Figure 11.**
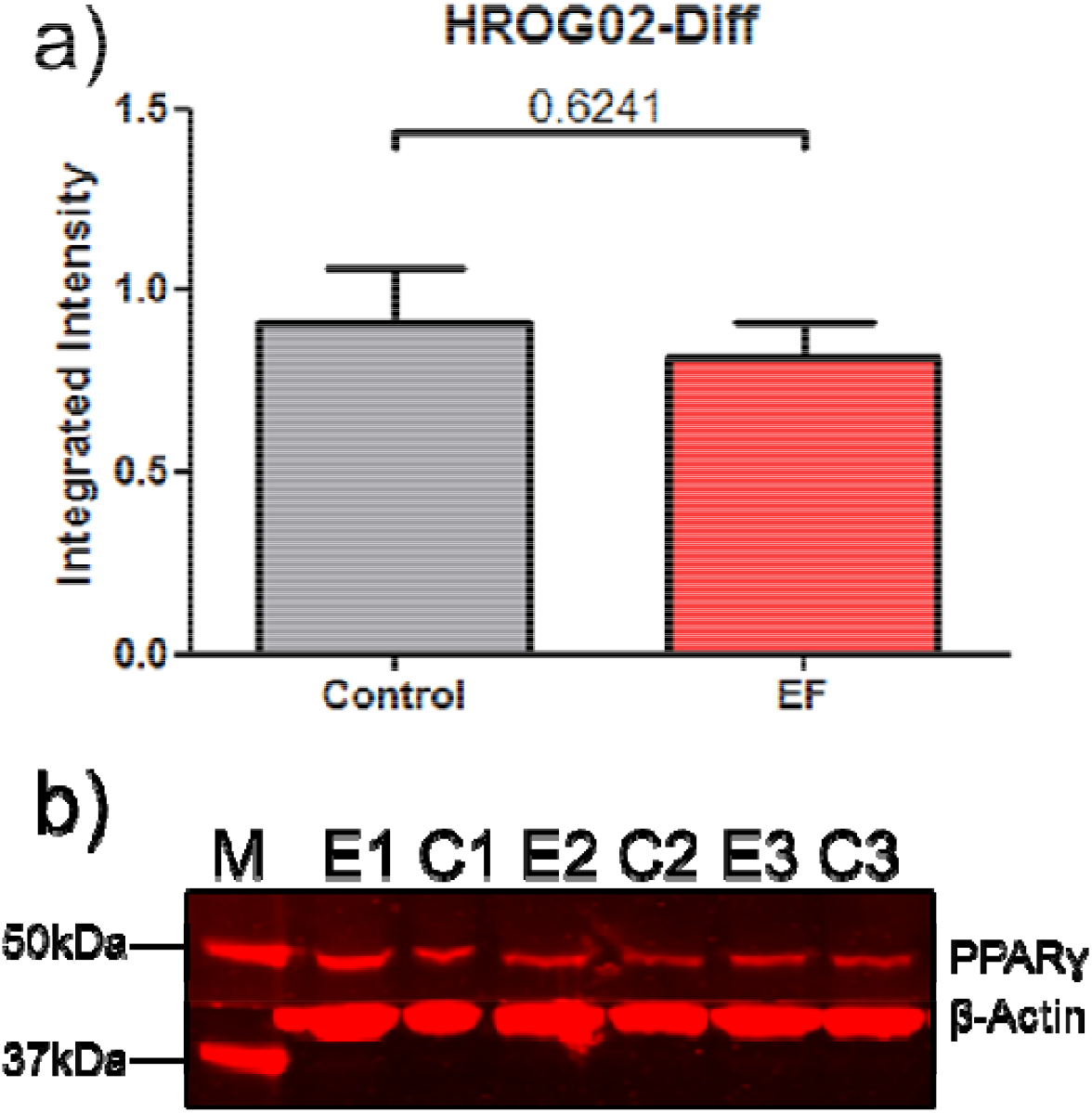
Effects of electric fields on PPARγ expression in HROG02-Diff cells. (a) shows that a one hour application on a 200mV/mm field did not cause a significant change in PPARγ protein levels when compared to β-actin expression. (b) is the image used in the Western blot analysis; M indicates the protein ladder, E indicates samples obtained from cells which were subjected to the electric field, C indicates control samples.

**Figure 12.**
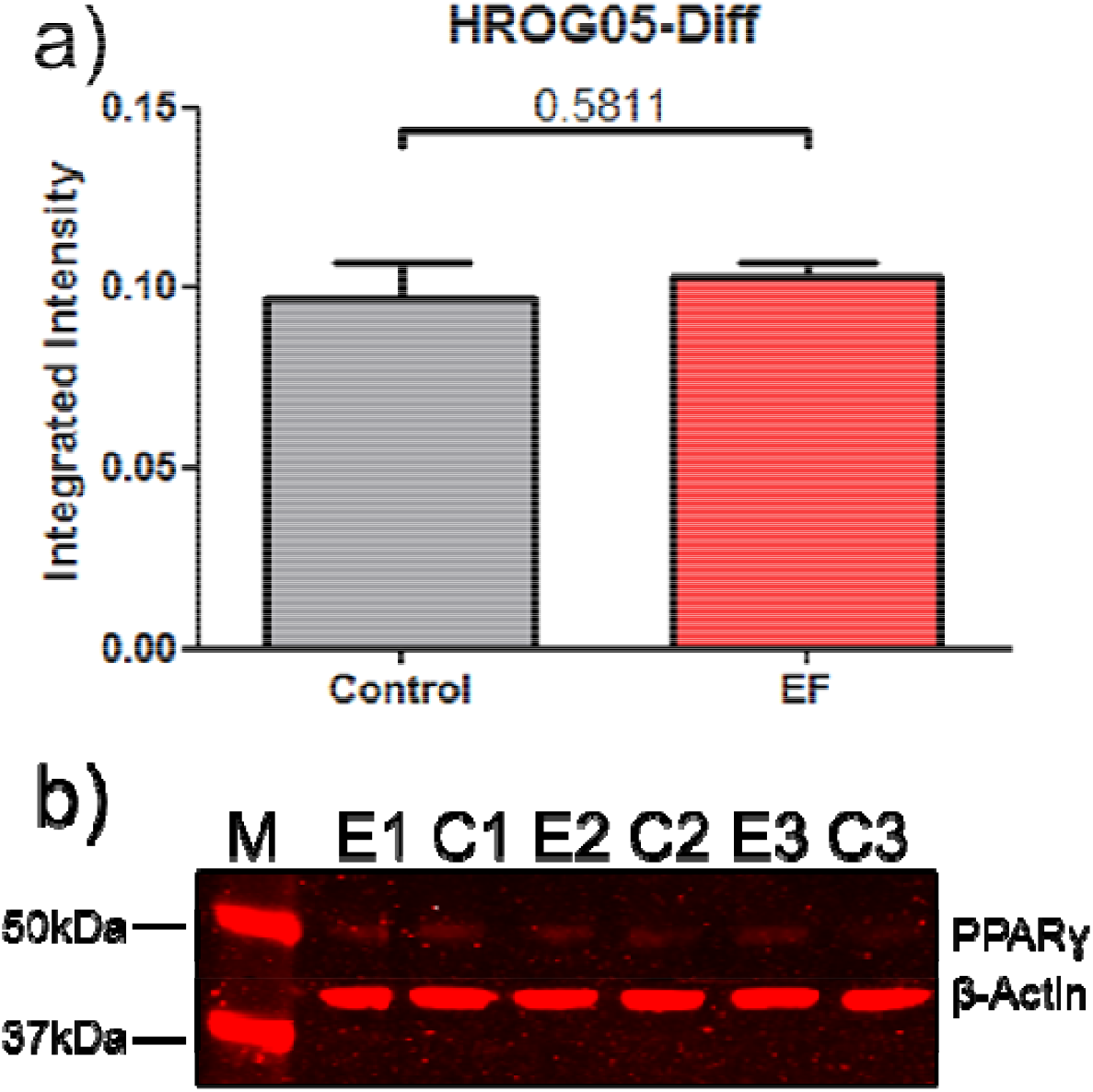
Effects of electric fields on PPARγ expression in HROG05-Diff cells. (a) shows that a one hour application on a 200mV/mm field did not cause a significant change in PPARγ protein levels when compared to β-actin expression. (b) is the image used in the Western blot analysis; M indicates the protein ladder, E indicates samples obtained from cells which were subjected to the electric field, C indicates control samples.

**Figure 13.**
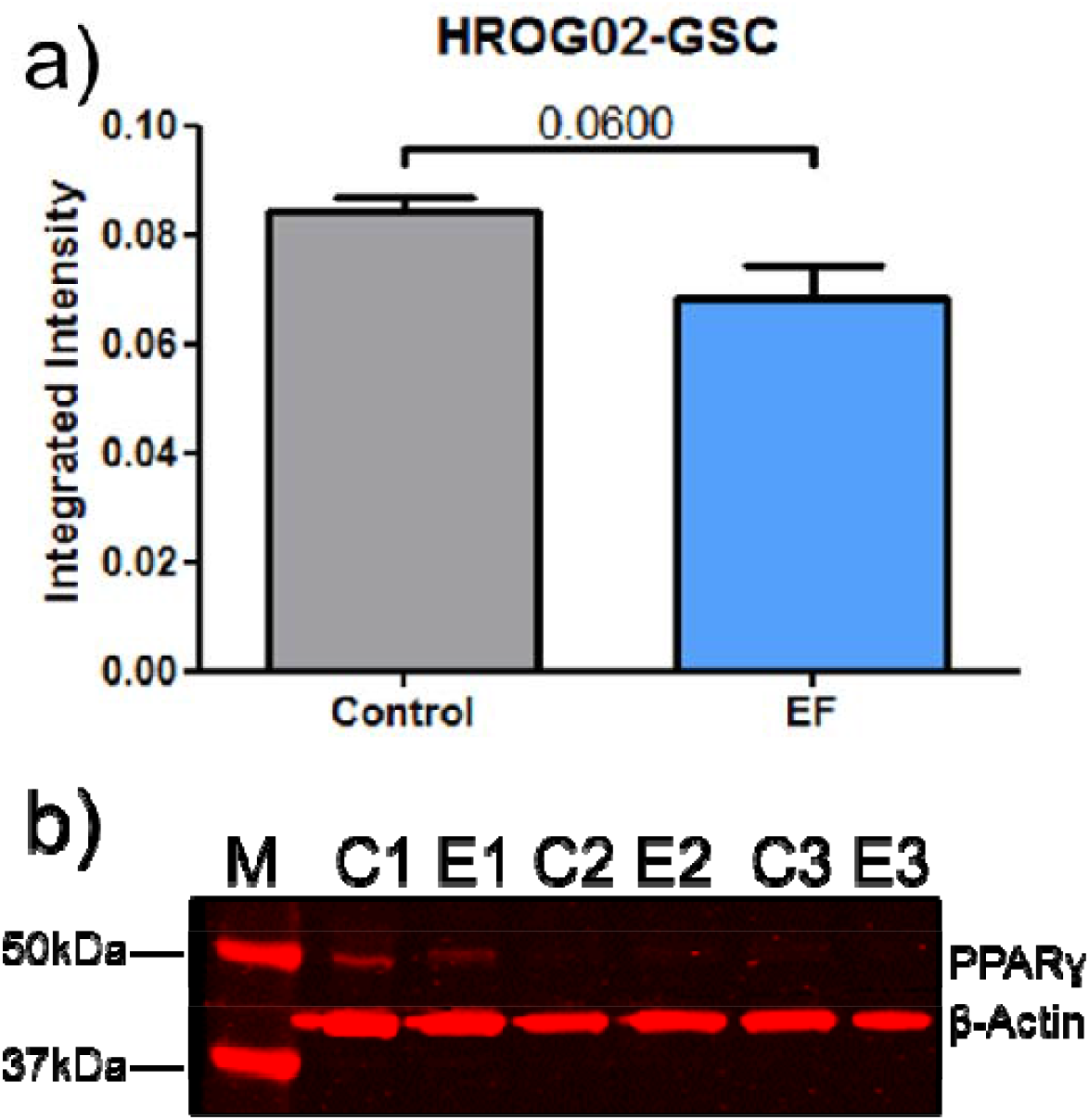
Effects of electric fields on PPARγ expression in HROG02-GSC cells. (a) shows that a one hour application on a 200mV/mm field did not cause a significant change in PPARγ protein levels when compared to β-actin expression. (b) is the image used in the Western blot analysis; M indicates the protein ladder, E indicates samples obtained from cells which were subjected to the electric field, C indicates control samples.

**Figure 14.**
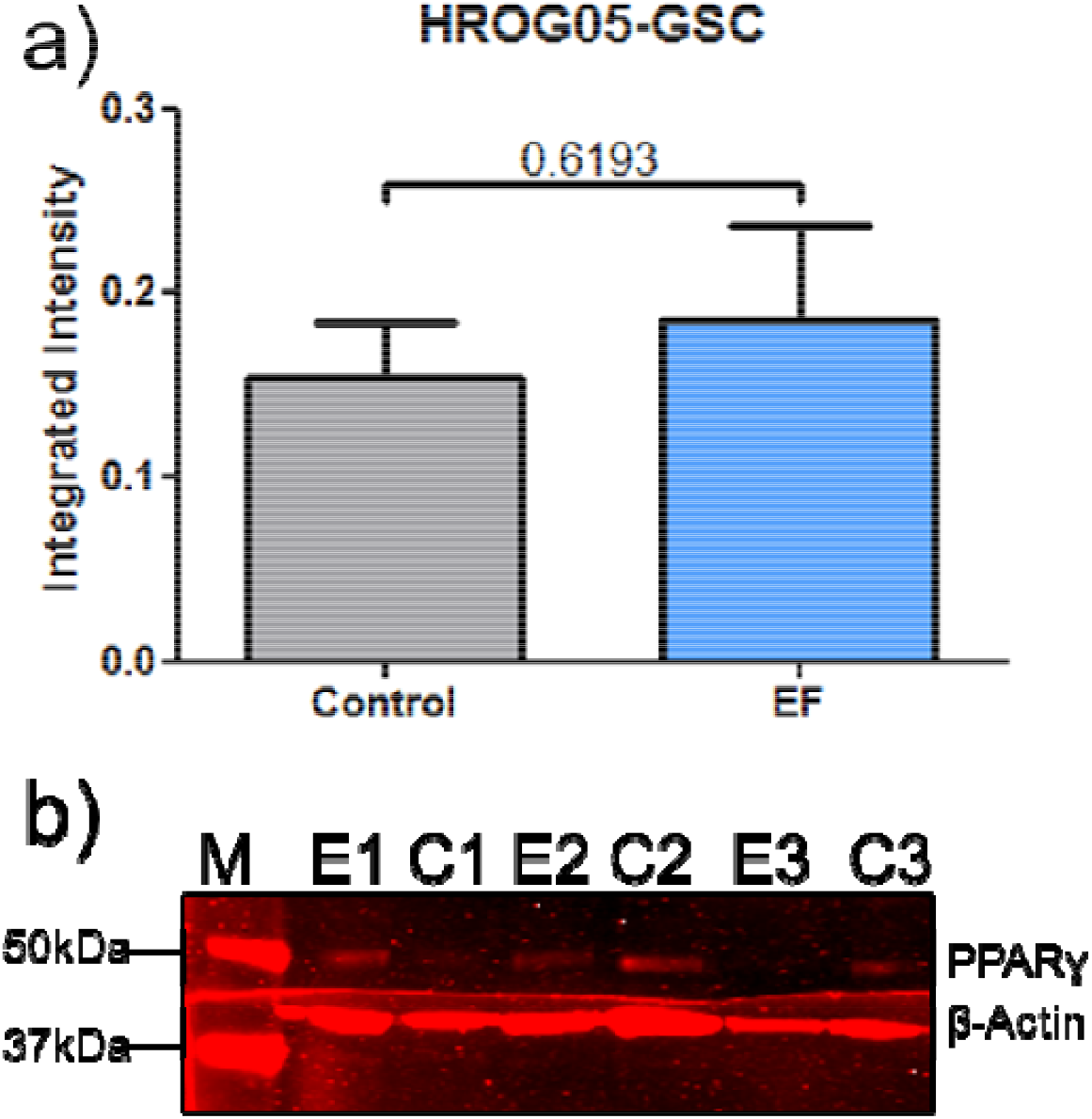
Effects of electric fields on PPARγ expression in HROG05-GSC cells. (a) shows that a one hour application on a 200mV/mm field did not cause a significant change in PPARγ protein levels when compared to β-actin expression. (b) is the image used in the Western blot analysis; M indicates the protein ladder, E indicates samples obtained from cells which were subjected to the electric field, C indicates control samples.

### Discussion

Herein we demonstrate for the first time that the application of an EF produces contrasting migratory responses in the differentiated and GSC-like states of primary GBM cell lines. Specifically, HROG02 and HROG05 cells migrated anodally in the differentiated state but reversed to cathodal migration in the de-differentiated, GSC-like state. Crucially this electrotaxis occurred at predefined physiological voltage gradients (50mV/mm)[20,21]. Our findings compliment recent evidence by Huang and colleagues (2016), who also demonstrate cathodal migration of 5 different patient derived brain tumour initiating cell lines in a 2-dimensional (2D) culture over poly-L-ornithine/laminin and in a 3D microfluidic device. However, Huang and colleagues were unable to recapitulate an opposing electrotactic response in the immortalised differentiated U87 GBM cell line, where no migratory response was elicited at 1V/cm, despite Li and colleagues reporting that U87MG cells migrate cathodally when exposed to 2V/cm [27,41]. Our observations indicate that GBM cells will migrate in opposite directions depending on their transition state, differentiated or stem-like, when exposed to an EF.

Similar contrasting electrotactic responses have been reported in other cancer types and are often related to metastatic potential. Prostate cancer cells, for example, with high and low metastatic potential migrate cathodally and anodally, respectively [42]. Further, a positive correlation between the metastatic potential and anodal migration has been reported in breast cancer cells [26]. This migratory phenomenon may therefore represent a malignant feature of cancer; opposing responses of subpopulations of cancer cells induces selective migration away from the tumour bulk and less predictable infiltration. This may explain the diffuse infiltration typically seen in GBM. In theory, if an electrical gradient existed between the tumour and the surrounding brain parenchyma, a subpopulation of either differentiated or GSC GBM cells would migrate away from the tumour depending on the electrical gradient polarity. Therefore, if diffuse GBM cell migration is time dependent, preventing or reversing this process may be more feasible earlier in the disease course, however this concept has yet to be fully investigated and may not be clinically feasible based on the requirements for medical screening tests [43]. However, the Food and Drug Administration approved in 2005 Novocure’s Tumour Treatment Fields (renamed recently to Optune), which utilises alternating current across the cranium in recurrent post-surgical GBM in combination with chemotherapy [44]. As such, the use of a direct current EF to manipulate GBM cell migration has yet to be optimised for clinical use.

Neural stem cells are known to be prone to transformation into GSCs [45] and have been demonstrated to reliably migrate in a cathodal direction in response to EFs in vitro and in vivo. [46,47,27]). In contrast, we and others find that GSCs migrate anodally, suggesting a pathological mechanism underlying this change in response [27]. Previous evidence implicates PI3K in electrotaxis, where cytoskeletal rearrangements ultimately lead to directed migration via Src/PI3K/Akt and MAPK signalling pathways [28,26,20,48]. Huang and colleagues demonstrate that chemical inhibition of PI3K with LY294002 decrease brain tumour initiating cells directed migration, where inhibition of Erk and each of its activating growth factors had no effect on electrotaxis [27]. Recently, Lyon and colleagues identified putative electrotactic signalling pathways with transciptomic analyses in differentiated U87MG spheroid aggregates, and subsequently finding PI3K, Akt, mTOR, ErbB2, ErbB3, Src/Abl, but not MEK, HGF/VEGF, ROCK1/2 inhibition attenuates cathodal migration [49]. However, PI3K inhibition with LY294002 did not affect cathodal migration, although BEZ235 abolished directed migration, which may be non-specifically accrued to its duel activity on PI3K/mTOR and more efficient downregulation of Akt [49,50]. Taken together, PI3K does not appear to be an efficient target to prevent directed cell migration in differentiated GBM cells with in vitro efficacy proven in GSCs but not differentiated GBM cells.

Based on findings that PPARγ agonists suppress Akt in both differentiated and GSC cells through PPARγ activation [31,51], we utilised pioglitazone to determine whether EF migration is affected by Akt downregulation by PPARγ. We found that pioglitazone treatment significantly decreased the directedness of electrotaxis of primary GBM cell lines in both differentiated and de-differentiated states. This inhibitory effect was diminished by the PPARγ antagonist GW9662, directly implicating PPARγ activation in suppression of EF guided migration of primary GBM cell lines. PPARγ activation inhibited electrotaxis in both differentiated and de-differentiated phenotypes implies a common migratory pathway downstream of PPARγ, such as the PI3K/Akt pathway. This provides early evidence that inhibition of GBM directed cell migration reliant on electrotaxis may be targeted with drug therapies. Further investigation including transcriptomics, selective chemical inhibition and in vivo experiments would be necessary to validate this. We present promising findings that a single drug can simultaneously inhibit differentiated and stem cell populations of the same primary GBM cells. As such PPARγ agonists may have a further clinical role in preventing GBM recurrence in addition to anti-proliferative, pro-differentiation of GSC and pro-apoptotic effects reported previously [52]. Such findings need to be recapitulated in animal models of GBM to determine whether tumour infiltration can be reversed. Such a strategy holds clinical importance as patients with GBM present late in the disease, making cancer cell migration reversal using electric fields a more realistic approach as opposed to a containment strategy such as chondroitin sulfate proteoglycans modulation, which would not be able to target satellite GBM cells [53].

Physiological EFs have been demonstrated in the mammalian brain [18] and such EF’s have been shown to occur in many other cancer types including ovarian, leukaemic, breast, cervical and prostate cancer cells. In these instances, depolarisation is positively correlated with the rate of proliferation [54]. Recently, Jun-Feng and colleagues demonstrate that human neural stem cells implanted in the rat brain can be electrically guided along the rostral migratory stream [47]. Not only does this have implications for neural regeneration but also for GBM treatment, where EF therapy may be useful as a treatment adjunct to prevent tumour recurrence. Interestingly, voltage-gated Na^+^ channels have been implicated in tumour-parenchyma EF generation [54] and GBMs demonstrate an association with metastatic potential and Na^+^ influx [55]. Furthermore, epileptiform hyperexcitability has been demonstrated at the interface between GBM and normal brain tissue, peri-tumour, that is mediated by glutamate excess [56]. While tumour epilepsy does not occur universally in patients with GBM, EF generation at the peri-tumour caused by increased extracellular glutamate may contribute to driving cell migration away from the tumour [57].

The generation of a pathological EF between GBM and surrounding brain parenchyma may be dependent on high rates of cellular proliferation and deregulated sodium transport. This would produce a relatively depolarised tumour bulk which would act as an anode whilst the surrounding brain tissue acts as a cathode. The cathodal migration of GSC-like primary GBM cell lines that we report here demonstrates a pro-migratory mechanism of highly tumourigenic GBM cells away from the tumour bulk. It is unknown how the presence of raised extracellular glutamate may affect this electrochemical gradient and requires further investigation to clarify the underlying mechanisms.

Further investigation including in vivo models and clinical trials will be required to prove whether electrotaxis of GBM and GSCs can be targeted chemically and/or surgically to add to the current armamentarium in brain tumour treatment.

### Conclusions

We demonstrate <u>for the first</u> time that differentiated and stem cell state primary GBMs derived from the same cell populations migrate in opposing directions in an applied EF. The directedness of both electrotaxis behaviours irrespective of polarity was reduced markedly by the PPARγ agonist pioglitazone. Regardless of the direction of EF present, cancer cells appear to migrate away from the tumour bulk. Further research into the generation and characteristics of tumour-brain EFs would provide invaluable insights into the *vivo* pathology. Pathological EF disruption may represent a novel target to prevent tumour infiltration. Chemical or surgical interventions may be developed to take advantage of this potential susceptibility in the future.

## Acknowledgements

We thank Dr Filipa Cunha, Dr Lin Cao and Dr Jin Pu for technical support in performing electroxis experiments. We thank Professor Michael Linnebacher for gifting primary brain tumour cell lines.

## Authors’ contributions

Conceptulisation, J.C., C.D.M.; Methodology, J.C., C.D.M.; Validation J.C. H.C., M.P.; Formal Analysis, H.P., M.P.; Investigation, J.C., H.P., M.P.; Data Curation, H.P., M.P., J.C.; Manuscript draft, H.P., M.P., J.C.; Manuscript final review, All; Supervision, C.D.M, J.C., B.L.; Funding, J.C., C.D.M.

## Ethics approval and consent to participate

Ethical approvals were obtained from the National Research Ethics Service – North of Scotland, REC reference: 14/NS/0015, Protocol No.: 3/076/13, IRAS project ID: 138989. Informed written and verbal consent was obtained from patients agreeing to participate in the study using the protocol approved by the North of Scotland Research Ethics Committee.

## Consent for publication

Not applicable.

## Data availability

All data generated or analysed during this study are included in this published article and its supplementary information files.

## References

1. Louis DN, Perry A, Reifenberger G, von Deimling A, Figarella-Branger D, Cavenee WK, Ohgaki H, Wiestler OD, Kleihues P, Ellison DW (2016) The 2016 World Health Organization Classification of Tumors. Acta Neuropathologica 131:803–820

2. Ostrom Qt, Gittleman H, Liao P, Rouse C, Chen Y, Dowling J, Wolinksy Y, Kruchko C, Barnholtz-Sloan J (2014) CBTRUS Statistical Report: Primary Brain and Central Nervous System Tumors Diagnosed in the United States in 2007-2011. Neuro-oncology 16:Suppl: iv1–iv63

3. Thakkar JP, Dolecek TA, Horbinski C, Ostrom QT, Lightner DD, Bamholtz-Sloan JS, Villano JL (2014) Epidemiologic and Molecular Prognostic Review of Glioblastoma. Cancer Epidemiology, Biomarkers & Prevention 23 (10):1985–1996

4. Stupp R, Hegi ME, Mason WP, van den Bent MJ, Taphoorn MJB, Janzer RC, Ludwin SK, Allgeier A, Fisher BA, Belanger K, Hau P, Brandes AA, Gijtenbeek J, Marosi C, Vecht CJ, Mokhtari K, Wesseling P, Villa S, Eisenhauer E, Gorlia T, Weller M, Lacombe D, Cairncross JG, Mirimanoff R (2009) Eff ects of radiotherapy with concomitant and adjuvant temozolomide versus radiotherapy alone on survival in glioblastoma in a randomised phase III study: 5-year analysis of the EORTC-NCIC trial. Lancet Oncology 10:459–466

5. Xie Q, Mittal S, Berens ME (2014) Targeting adaptive glioblastoma: an overview of proliferation and invasion. Neuro-Oncology 16 (12):1575–1584

6. Ignatova TN, Kukekov VG, Leywell ED, Suslov ON, Vrionis FD, Steindler DA (2002) Human Cortical Glial Tumors Contain Neural Stem-Like Cells Expressing Astroglial and Neuronal Markers In Vitro. GLIA 39:193–206

7. Singh SK, Clarke ID, Terasaki M, Bon VE, Hawkins C, Squire J, Dirks PB (2003) Identification of a Cancer Stem Cell in Human Brain Tumors. Cancer Research 63:5821–5828

8. Hemmati HD, Nakano I, Lazareff JA, Masterman-Smith M, Geschwind DH, Bronner-Fraser M, Kornblum HI (2003) Cancerous stem cells can arise from pediatric brain tumors. PNAS 100 (25):15178–15183

9. Galli R, Binda E, Orfanelli U, Cipelletti B, Gritti A, De Vitis S, Fiocco R, Foroni C, Dimeco F, Vescovi A (2004) Isolation and Characterization of Tumorigenic, Stem-like Neural Precursors from Human Glioblastoma. Cancer Research 64:7011–7021

10. Lathia JD, Mack SC, Mulkearns-Hubert EE, Valentim CLL, Rich JN (2015) Cancer stem cells in glioblastoma. Genes and Development 29:1203–1217

11. Shin JH, Lee YS, Hong Y, Kang CS (2013) Correlation between the prognostic value and the expression of the stem cell marker CD133 and isocitrate dehydrogenase1 in glioblastomas. Journal of Neurooncology 115:333–341

12. Pallini R, Ricci-Vitiani L, Banna GL, Signore M, Lombardi D, Todaro M, Stassi G, Martini M, Maira G, Larocca LM, De Maria R (2008) Cancer Stem Cell Analysis and Clinical Outcome in Patients with GlioblastomaMultiforme. Clinical Cancer Research 14 (24):8205–8212

13. Demuth T, Berens ME (2004) Molecular Mechanisms of Glioma Cell Migration and Invasion. Journal of Neuro-Oncology 70

14. Lefranc F, Brotchi J, Kiss R (2005) Possible Future Issues in the Treatment of Glioblastomas: Special Emphasis on Cell Migration and the Resistance of Migrating Glioblastoma Cells to Apoptosis. Journal of Clinical Oncology 23:2411–2422

15. Paw I, Carpenter RC, Watabe K, Debinski W, Lo H (2015) Mechanisms regulating glioma invasion. Cancer Letters 362:1–7

16. Ariza CA, Fleury AT, Tormos CJ, Petruk V, Chawla S, Oh J, Sakaguchi DS, Mallapragada SK (2010) The Influence of Electric Fields on Hippocampal Neural Progenitor Cells. Stem Cell Reviews and Reports 6:585–600

17. Arocena M, Zhao M, Collinson JM, Song B (2010) A Time-Lapse and Quantitative Modelling Analysis of Neural Stem Cell Motion in the Absence of Directional Cues and in Electric Fields. Journal of Neuroscience Research 88:3267–3274

18. Cao L, Wei D, Reid B, Zhao S, Pu J, Pan T, Yamoah E, Zhao M (2013) Endogenous Electric Currents Might Guide Rostral Migration of Neuroblasts. EMBO Reports 14:184–190

19. Li L, El-Hayek YH, Liu B, Chen Y, Gomez E, Wu X, Ning K, Li L, Chang N, Zhang L, Wang Z, Hu X, Wan Q (2008) Direct-Current Electrical Field Guides Neuronal Stem/Progenitor Cell Migration. Stem Cells 26:2193–2200

20. Meng X, Arocena M, Penninger J, Gage FH, Zhao M, Song B (2011) PI3K mediated electrotaxis of embryonic and adult neural progenitor cells in the presence of growth factors. Experimental Neurology 227:210–217

21. Yao L, Shanley L, McCaig C, Zhao M (2008) Small Applied Electric Fields Guide Migration of Hippocampal Neurons. Journal of Cellular Physiology 216:527–535

22. Zhao H, Steiger A, Nohner M, Ye H (2015) Specific Intensity Direct Current (DC) Electric Field Improves Neural Stem Cell Migration and Enhances Differentiation towards βIII-Tubulin+ Neurons. PLOS ONE 10 (6):1–21

23. McCaig CD, Song B, Rajnicek AM (2009) Electrical Dimensions in Cell Science. Journal of Cell Science 122:4267–4276

24. Li F, Chen T, Hu S, Lin J, Hu R, Feng H (2013) Superoxide Mediates Direct Current Electric Field-Induced Directional Migration of Glioma Cells through the Activation of AKT and ERK. PLOS ONE 8 (4):1–11

25. Mycielska ME, Djamgoz MBA (2004) Cellular Mechanisms of Direct-Current Electric Field Effects: Galvanotaxis and Metastatic Disease. Journal of Cell Science 117

26. Pu J, McCaig CD, Cao L, Zhao Z, Segall JE, Zhao M (2007) EGF receptor signalling is essential for electric-field directed migration of breast cancer cells. Journal of Cell Science 120:3395–3403

27. Huang Y, Hoffmann G, Wheeler B, Schiapparelli P, Quinones-Hinojosa A, Searson P (2016) Cellular microenvironment modulates the galvanotaxis of brain tumor initiating cells. Scientific Reports 6 (21583)

28. Zhao M, Song B, Pu J, Wada T, Reid B, Tai G, Wng F, Guo A, Walczysko P, Gu Y, Takehiko S, Suzuki A, Forrester JV, Bourne HR, Devreotes PN, McCaig CD, Penninger JM (2006) Electrical signals control wound healing through phosphatidylinositol-3-OH kinase-γ and PTEN. Nature 442:457–460

29. Zhao M, Pu J, Forrester JV, McCaig CD (2002) Zhao, M., Pu, Membrane lipids, EGF receptors and intracellular signals co-localize and are polarized in epithelial cells moving directionally in a physiological electric field. Federation of American Societies for Experimental Biology 16 (8):857–859

30. Mezey G, Treszl A, Schally AV, Block NL, Vízkeleti L, Juhász A, Nagy J, Balázs M (2014) Prognosis in human glioblastoma based on expression of ligand growth hormone-releasing hormone, pituitary-type growth hormone-releasing hormone receptor, its splicing variant receptors, EGF receptor and PTEN genes. Journal of Cancer Research and Clinical Oncology 140:1641–1649

31. Ching J, Amiridis S, Stylli SS, Bjorksten AR, Kountouri N, Zheng T, Paradiso L, Luwor RB, Morokoff AP, O’Brien TJ, Kaye AH (2015) The peroxisome proliferator activated receptor gamma agonist pioglitazone increases functional expression of the glutamate transporter excitatory amino acid transporter 2 (EAAT2) in human glioblastoma cells. Oncotarget:1–14

32. Patel L, Pass I, Coxen P, Downes CP, Smith SA, Macphee CH (2001) Tumor suppressor and anti-inflammatory actions of PPARg agonists are mediated via upregulation of PTEN. Current Biology 11 (10):765–768

33. Cimini A, Cristiano L, Colafarina S, Benedetti E, Di Loreto S, Festucciar C, Amicarelli F, Canuto RA, Ceru MP (2005) PPARc-dependent effects of conjugated linoleic acid on the human glioblastoma cell line (ADF). International Journal of Cancer 117:923–933

34. Zang C, Wachter M, Liu H, Posch MG, Fenner MH, Stadelmann C, von Deimling A, Possinger K, Black KL, Koeffler HP, Elstner E (2003) Ligands for PPARγ and RAR cause induction of growth inhibition and apoptosis in human glioblastomas. Journal of Neuro-Oncology 65:107–118

35. Cilibrasi C, Butta V, Riva G, Bentivegna A (2016) Pioglitazone Effect on Glioma Stem Cell Lines: Really a Promising Drug Therapy for Glioblastoma? PPAR Research 2016 (7175067):1–8

36. Fael Al-Mayhani TM, Ball SLR, Zhao J, Fawcett J, Ichmura K, Collins PV, Watts C (2009) An efficient method for derivation and propagation of glioblastoma cell lines that conserves the molecular [rofile of their original tumours. Journal of Neuroscience Methods: 192–199

37. Pollard SM, Yoshikawa K, Clarke ID, Danovi D, Stricker S, Russell R, Bayani J, Head R, Lee M, Bernstein M, Squire JA, Smith A, Dirk P (2009) Glioma Stem Cell Lines Expanded in Adherent Culture Have Tumor-Specific Phenotypes and Are Suitable for Chemical and Genetic Screens. Cell:568–580

38. Chearwae W, Bright JJ (2008) PPARg agonists inhibit growth and expansion of CD133+ brain tumour stem cells. British Journal of Cancer 99:2044–2053

39. Song B, Gu Y, Pu J, Reid B, Zhao Z, Zhao M (2007) Application of direct current electric fields to cells and tissues in vitro and modulation of wound electric field in vivo. Nature Protocols 2 (6):1479–1489

40. Vik-Mo EO, Sandberg C, Olstom H, Varghese M, Brandal P, Ramm-Pettersen J, Murrell W, Langmoen IA (2010) Brain tumor stem cells maintain overall phenotype and tumorigenicity after in vitro culturing in serum-free conditions. Neuro-Oncology 12 (12):1220–1230

41. Li F, Chen T, Hu S, Lin J, Hu R, Feng H (2013) Superoxide Mediates Direct Current Electric Field-Induced Directional Migration of Glioma Cells through the Activation of AKT and ERK vol 8. PLoS ONE,

42. Djamgoz M, Mycielska M, Madeja Z, Fraser SP, Korohoda W (2001) Directional movement of rat prostate cancer cells in direct-current electric field: involvement of voltagegated Na+ channel activity.. Journal of Cell Science 114 (14):2697–2705

43. Pinsky PF (2015) Principles of Cancer Screening. Surg Clin North Am 95 (5):953–966. doi:10.1016/j.suc.2015.05.009

44. Fabian D, Guillermo Prieto Eibl MDP, Alnahhas I, Sebastian N, Giglio P, Puduvalli V, Gonzalez J, Palmer JD (2019) Treatment of Glioblastoma (GBM) with the Addition of Tumor-Treating Fields (TTF): A Review. Cancers (Basel) 11 (2). doi:10.3390/cancers11020174

45. Vescovi AL, Galli R, Reynolds BA (2006) Brain tumour stem cells. Nat Rev Cancer 6 (6):425–436. doi:10.1038/nrc1889

46. Feng JF, Liu J, Zhang XZ, Zhang L, Jiang JY, Nolta J, Zhao M (2012) Guided migration of neural stem cells derived from human embryonic stem cells by an electric field. Stem Cells 30 (2):349–355. doi:10.1002/stem.779

47. Feng JF, Liu J, Zhang L, Jiang JY, Russell M, Lyeth BG, Nolta JA, Zhao M (2017) Electrical Guidance of Human Stem Cells in the Rat Brain. Stem Cell Reports 9 (1):177–189. doi:10.1016/j.stemcr.2017.05.035

48. Sun Y, Do H, Gao J, Zhao R, Zhao M, Mogilner A (2013) Keratocyte fragments and cells utilize competing pathways to move in opposite directions in an electric field. Curr Biol 23 (7):569–574. doi:10.1016/j.cub.2013.02.026

49. Lyon JG, Carroll SL, Mokarram N, Bellamkonda RV (2019) Electrotaxis of Glioblastoma and Medulloblastoma Spheroidal Aggregates. Sci Rep 9 (1):5309. doi:10.1038/s41598-019-41505-6

50. Maira SM, Stauffer F, Brueggen J, Furet P, Schnell C, Fritsch C, Brachmann S, Chène P, De Pover A, Schoemaker K, Fabbro D, Gabriel D, Simonen M, Murphy L, Finan P, Sellers W, García-Echeverría C (2008) Identification and characterization of NVP-BEZ235, a new orally available dual phosphatidylinositol 3-kinase/mammalian target of rapamycin inhibitor with potent in vivo antitumor activity. Mol Cancer Ther 7 (7):1851–1863. doi:10.1158/1535-7163.MCT-08-0017

51. Im CN (2017) Combination Treatment with PPAR. Biomed Res Int 2017:5832824. doi:10.1155/2017/5832824

52. Ellis HP, Kurian KM (2014) Biological Rationale for the Use of PPARγ Agonists in Glioblastoma. Front Oncol 4:52. doi:10.3389/fonc.2014.00052

53. Silver DJ, Siebzehnrubl FA, Schildts MJ, Yachnis AT, Smith GM, Smith AA, Scheffler B, Reynolds BA, Silver J, Steindler DA (2013) Chondroitin sulfate proteoglycans potently inhibit invasion and serve as a central organizer of the brain tumor microenvironment. J Neurosci 33 (39):15603–15617. doi:10.1523/JNEUROSCI.3004-12.2013

54. Yang M, Brackenbury J (2013) Membrane Potential and Cancer Progression. Frontiers in Physiology 4:1–10

55. Berdiev BK, Xia J, McLean LA, Markert JM, Gillespie GY, Mapstone TB, Naren AP, Jovov B, Bubien JK, Ji H, Fuller CM, Kirk KL, Benos DJ (2003) Acid-sensing Ion Channels in Malignant Gliomas. The Journal of Biological Chemistry 278 (17):15023–15034

56. Buckingham SC, Campbell SL, Haas BR, Montana V, Robel S, Ogunrinu T, Sontheimer H (2011) Glutamate release by primary brain tumors induces epileptic activity. Nat Med 17 (10):1269–1274. doi:10.1038/nm.2453

57. Yuen TI, Morokoff AP, Bjorksten A, D’Abaco G, Paradiso L, Finch S, Wong D, Reid CA, Powell KL, Drummond KJ, Rosenthal MA, Kaye AH, O’Brien TJ (2012) Glutamate is associated with a higher risk of seizures in patients with gliomas. Neurology 79 (9):883–889. doi:10.1212/WNL.0b013e318266fa89

